# The *Arabidopsis* 14-3-3 proteins stabilize BRASSINOSTEROID INSENSITIVE1 at the plasma membrane

**DOI:** 10.1101/2024.12.19.629515

**Authors:** Luyao Li, Chengzhu Yin, Chunyan Yan, Zi-Liang Hu, Isabelle Vanhoutte, Yan Li, Bingbing Tang, Guorong Xiao, Xinyuan Feng, Wenhui Cao, Yong Wang, Dexian Luo, Eugenia Russinova, Derui Liu

## Abstract

Brassinosteroid (BR) hormones regulate various physiological and developmental processes in plants. BR signaling is primarily influenced by the plasma membrane abundance of the BR receptor BR INSENSITIVE1 (BRI1), a process regulated by ubiquitination, endocytosis, and protein degradation. Despite extensive research, only a few negative regulators of BRI1 internalization and ubiquitination have been identified. In this study, we show that the conserved eukaryotic regulatory proteins 14-3-3 directly interact with BRI1 at Threonine 872 (T872) within its juxtamembrane domain. Furthermore, phosphorylation at Serine 858 (S858) in BRI1’s juxtamembrane domain enhances T872 phosphorylation, facilitating 14-3-3 protein binding. Consequently, by inhibiting BRI1 ubiquitination without affecting its kinase activity or BAK1 interaction, 14-3-3 binding increased BRI1 plasma membrane abundance and enhanced BR signaling. Both non-epsilon and epsilon isoforms of 14-3-3 proteins contribute to the regulation of BRI1 and, consequently, to plant responsiveness to BRs. Our results revealed a previously undescribed function of 14-3-3 proteins in regulating BRI1 stability.

**One Sentence Summary:** 14-3-3s stabilize BRI1 by antagonizing its ubiquitination

## Introduction

Brassinosteroids (BRs) are steroidal phytohormones that are essential for plant growth, development, and response to abiotic and biotic stresses (Nolan et al., 2020). BR signaling is initiated by the binding of the hormone to its plasma membrane (PM)-localized cognate receptor kinase (RK), BR INSENSITIVE1 (BRI1) (Friedrichsen et al., 2000; He et al., 2000) and its homologs, BRI1-LIKE1 (BRL1) and BRL3 (Cano-Delgado et al., 2004; Kinoshita et al., 2005). In the absence of BRs, the inhibitory protein BRI1 KINASE INHIBITOR1 (BKI1) (Wang and Chory, 2006; Jaillais et al., 2011; Jiang et al., 2015) primarily maintains BRI1 in an inactive state. However, upon binding to BRs, BRI1 heterodimerizes with its co-receptor BRI1-ASSOCIATED KINASE1 (BAK1), also known as SOMATIC EMBRYOGENESIS RECEPTOR KINASE3 (SERK3) and its homologs SERK1 and SERK4. During this process, BRI1 and BAK1 auto- and trans-phosphorylate to form fully activated receptors (Wang et al., 2008). BRs binding to BRI1 also triggers dissociation of its negative regulators, BKI1 and BOTRYTIS-INDUCED KINASE1 (BIK1) (Wang et al., 2011; Lin et al., 2013). The BR signal is transduced from the PM to the nucleus through a sequential phosphorylation and dephosphorylation cascade to ultimately dephosphorylate and activate the master transcription factors, BRASSINAZOLE-RESISTANT1 (BZR1) and BRI1-EMS-SUPPRESSOR1 (BES1)/BZR2, which further control BR-responsive gene expression (Nolan et al., 2020).

The key determinant of BR signaling is the abundance of BRI1 in the PM, which is controlled by its ubiquitination, endocytosis and degradation (Di Rubbo et al., 2013; Gadeyne et al., 2014; Martins et al., 2015; Zhou et al., 2018; Liu et al., 2020). Ligand-dependent activation and phosphorylation of BRI1 promotes its association with the plant U-box (PUB) E3 ubiquitin ligases PUB12 and PUB13, which in turn directly ubiquitinate BRI1. Importantly, PUB13 is phosphorylated by BRI1 on a specific residue, enhancing their association. The ubiquitinated BRI1 is subsequently endocytosed and sorted for vacuolar degradation (Martins et al., 2015; Zhou et al., 2018). The ubiquitination of BRI1 can be alleviated by deubiquitinating enzymes UBP12 and UBP13, thereby stabilizing it (Luo et al., 2022). In addition to ubiquitination-dependent endocytosis, BRI1 regulates its PM pool through clathrin-mediated endocytosis (Di Rubbo et al., 2013; Gadeyne et al., 2014; Liu et al., 2020), which involves the adaptor protein complex-2 (AP-2) and TPLATE complex (TPC), as well as clathrin-independent endocytosis (Wang et al., 2015), occurring with or without BRs binding (Geldner et al., 2007; Claus et al., 2023).

14-3-3 proteins (hereafter 14-3-3s), also known as general regulatory factors (GRFs), are a family of regulatory proteins that are highly conserved in eukaryotes (Darling et al., 2005). Although they do not confer enzymatic activity, they function as dimers, with each monomer binding to phosphorylated (p) serine (S) or threonine (T) residues within their target proteins, making 14-3-3s also known as bridging proteins for protein-protein interactions. Three sequence-specific motifs, RSxpSxP, RSxxpSxP, and YpT, are the common phosphorylated peptides for 14-3-3 binding. However, 14-3-3s can also interact with phosphorylated proteins lacking these conserved motifs (Paul et al., 2012). Furthermore, although rare, phosphorylation-independent interactions with 14-3-3s have also been reported (Wang et al., 1999; Fuglsang et al., 2003).

The *Arabidopsis thaliana* (Arabidopsis) genome encodes 13 14-3-3 isoforms, which based on their sequence characteristics, are divided into two subfamilies: non-epsilon, including GRF1 to GRF8, and epsilon, including GRF9 to GRF13 (Denison et al., 2011). Although 14-3-3 isoforms share protein structural similarities and some functional redundancy, their targets and physiological functions may differ due to differences in spatiotemporal pattern of gene expression and subcellular localization (Alsterfjord et al., 2004; Paul et al., 2012; Pallucca et al., 2014; Keicher et al., 2017).

14-3-3s interact with numerous proteins to regulate their enzymatic and binding activity (Lambeck et al., 2010; Yang et al., 2019; Dong et al., 2023; Ma et al., 2023), subcellular localization (Gampala et al., 2007; Wang et al., 2011; Keicher et al., 2017; Sullivan et al., 2021; Fan et al., 2023), and turnover (Qi et al., 2022) in plant cells. In the absence or low concentration of BRs, 14-3-3s act as negative regulators by binding the phosphorylated BES1 and BZR1 in the cytoplasm, thereby decreasing their nuclear abundance (Bai et al., 2007; Gampala et al., 2007). However, in the presence of BRs, 14-3-3 proteins bind to BKI1, promoting its dissociation from BRI1 at the PM, which in turn facilitates the nuclear localization of BES1 and BZR1 (Jaillais et al., 2011; Wang et al., 2011; Jiang et al., 2015), thereby positioning 14-3-3s as positive regulators in BR signaling. The nuclear localization of BZR1 is also positively regulated by Receptor for Activated C Kinase 1 (RACK1), which interacts with BZR1 and further reduces its association with 14-3-3 proteins in the cytoplasm (Li et al., 2023).

High-throughput proteomics revealed that in Arabidopsis besides BES1, BZR1 and BKI1, the GRF2 isoform associates with several BR signaling components, including BRI1, BAK1, the phosphatase *bri1-*SUPPRESSOR1 (BSU1) (Chang et al., 2009), and the GSK3-like kinase BR-INSENSITIVE2 (BIN2) (Kim et al., 2023). In addition, 14-3-3s have been shown to interact directly with SERK1 (Rienties et al., 2005).

Although studies have hinted at an association between 14-3-3s and BRI1, the molecular mechanism of their interactions was only studied *in vitro* (Chae et al., 2016; Obergfell et al., 2024). Isothermal Titration Calorimetry (ITC) and site-directed mutagenesis of synthetic peptides were used to demonstrate that BRI1 strongly binds the non-epsilon 14-3-3 isoforms Chi (χ; GRF1), Omega (ω; GRF2), Psi (ψ; GRF3) Phi (φ; GRF4), and Upsilon (υ; GRF5) (Chae et al., 2016) and that 14-3-3s recognize sequence-specific motifs in BRI1, BKI1, and BZR1 (Obergfell et al., 2024). The slightly reduced sensitivity to brassinazole (BRZ), an inhibitor of BR biosynthesis, observed in the quadruple non-epsilon 14-3-3 mutants including *grf1,4,6,8*, *grf5,6,7,8*, and *grf1,4,5,7* reinforced the idea that 14-3-3s function mainly as negative regulators of BR signaling (Obergfell et al., 2024). In contrast, although the triple *grf3,4,10* mutant did not display BR-related phenotypes, it was able to reduce the hypersensitivity to BRs in *BRI1* overexpression lines (Lee et al., 2020). These findings imply that 14-3-3 proteins might play a role in BR perception via BRI1 interaction.

Here, we demonstrate that all ubiquitously expressed Arabidopsis 14-3-3 proteins directly bind to BRI1, possibly at the S858 and T872 residues within its juxtamembrane domain. Mutation of S858 and T872 to alanine led to increased ubiquitination, followed by enhanced endocytosis and vacuolar targeting resulting in BRI1 degradation. Additionally, both non-epsilon higher-order mutant and epsilon RNAi plants exhibited BRI1 instability. Together, our findings establish 14-3-3 proteins as positive regulators of BRI1 abundance at the PM.

## Results

### 14-3-3s bind directly BRI1

Given that some 14-3-3 isoforms have been shown to associate with BRI1 and to confer specificity (Chang et al., 2009; Chae et al., 2016; Obergfell et al., 2024), we first tested whether isoforms co-expressed with BRI1 can interact. Of all the isoforms, *GRF1* to *GRF11* are ubiquitously expressed, which are very similar to the expression pattern of *BRI1* (Keicher et al., 2017). In contrast, *GRF12* is only expressed in flowers, and the expression level of *GRF13* is strikingly low (Wilson et al., 2016; Keicher et al., 2017). Therefore, GRF1 to GRF11 were selected for further analysis.

To determine whether GRFs interact directly with BRI1, we performed *in vitro* glutathione S-transferase (GST) pull-down assays using the bacterially expressed GST-tagged BRI1-cytoplasmic domain (CD), namely GST-BRI1-CD, and HIS-tagged GRFs. GST-BRI1-CD pulled down almost all GRF isoforms tested. The interaction between BRI1-CD and GRF2, GRF5, GRF6, GRF8, GRF9, and GRF11 was the strongest, whereas the interaction between BRI1 and GRF3, GRF4, and GRF10 was moderate (Fig. 1A). In contrast, interactions with GRF1 and GRF7 were scarce, and no interactions were detected between GST (negative control) and each of HIS-GRF3, HIS-GRF6, and HIS-GRF9 (Fig. 1A). These data indicate that different 14-3-3 protein isoforms may exhibit altered binding affinity to BRI1.

**Figure 1.**
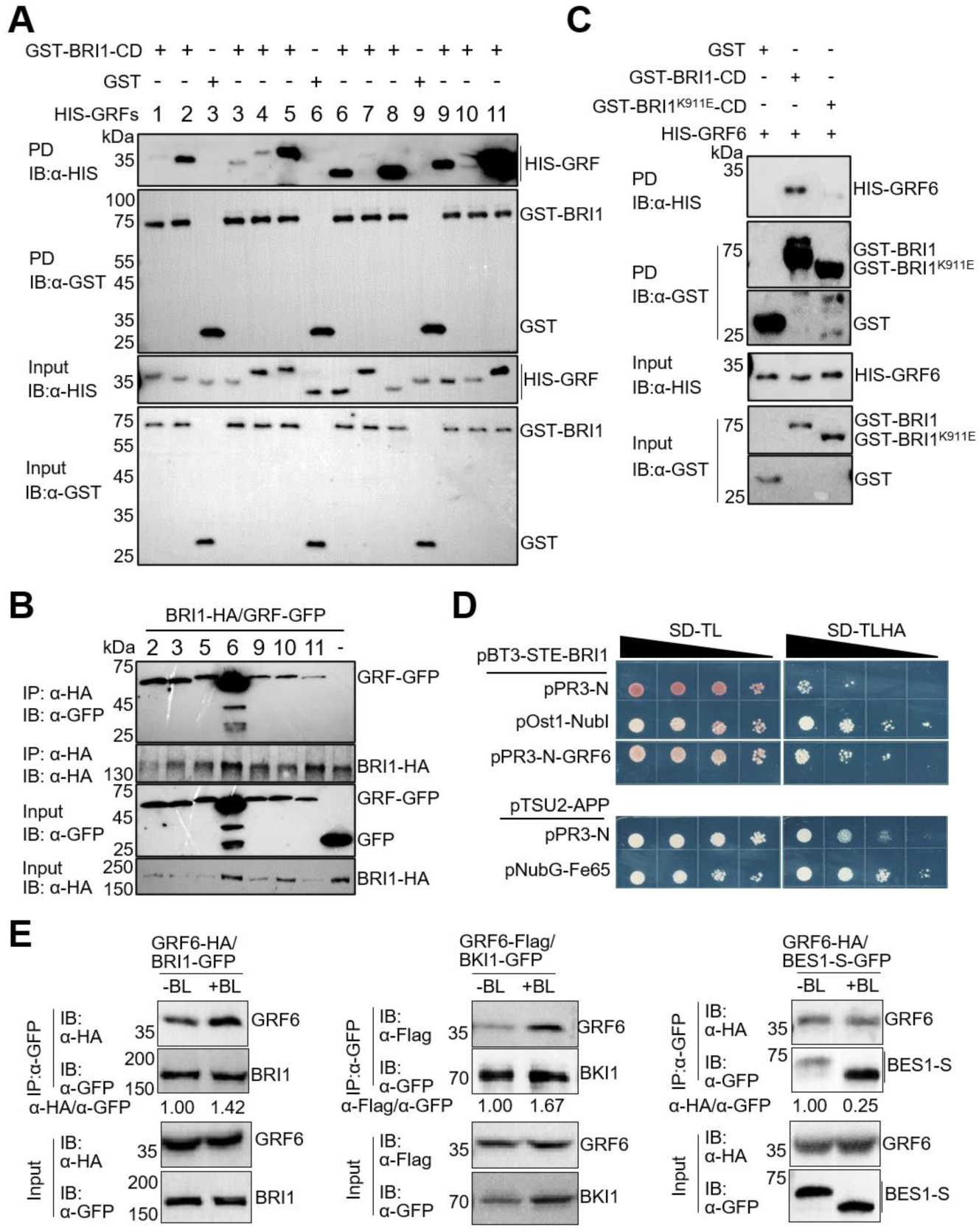
14-3-3s interact directly with BRI1. **A**) GRF1 to GRF11 interact with BRI1 *in vitro*. HIS-fused GRFs were incubated with GST or GST-fused BRI1-cytoplasmic domain (CD; GST-BRI1-CD). The beads were collected, washed, and immunoblotted (IB) with α-GST and α-HIS antibodies. Protein inputs were determined with α-GST and α-HIS antibodies. **B**) Association between BRI1 and GRFs *in planta*. *N. benthamiana* leaves co-expressing GFP-tagged GRFs or free GFP together with HA-tagged BRI1 were collected 48 h post-agroinfiltration. Co-immunoprecipitation (co-IP) assays using α-HA beads were done with proteins isolated from these samples. Proteins were subjected to immunoblot (IB) assay using α-HA and α-GFP antibodies. **C**) GRF6 associations with wild-type BRI1 and BRI1^K911E^ mutant determined by *in vitro* GST pull-down. HIS-fused GRF6 was incubated with free GST, GST-fused BRI1-CD, or BRI1^K911E^-CD. The beads were collected and washed, and the pull-down fractions were followed by IBs with α-GST and α-HIS. Protein inputs were determined by IB with α-GST and α-HIS antibodies. **D**) Mating based split-ubiquitin assay of the interaction between BRI1 and GRF6. *pTSU2-APP/pNubG-Fe65*, systemic positive control; *pTSU2-APP/pPR3N*, systemic negative control; *pBT3-STE-BRI1*/*pPR3N*, self-activation control; *pBT3-STE-BRI1/pOst1-NubI*, positive control for BRI1; and *pBT3-STE-BRI1/pPR3N-GRF6*, combination of interest. SD-TL, control medium (SD/-Trp-Leu); SD-TLHA, selective medium (SD/-Trp-Leu-His-Ade). Ten-fold serial dilutions of the saturated cultures were spotted onto plates. **E**) BL treatment enhanced BRI1 and GRF6 association. 5-day-old indicated lines were treated with 2 μM BRZ for 3 days, followed by 1 μM BL treatment (+BL) or continuously with 2 μM BRZ (-BL) for 1.5h. co-IP assays using α-GFP beads were done with proteins isolated from these samples. Proteins were subjected to IB using α-HA, α-GFP, and α-Flag antibodies. The blots were performed twice, and similar results were obtained.

To verify the association between BRI1 and GRFs *in planta*, co-immunoprecipitation (co-IP) assays were carried out by transient expressing HA-tagged BRI1 together with the GFP-tagged isoforms GRF2, GRF3, GRF5, GRF6, GRF9, GRF10, and GRF11 or free GFP in *Nicotiana benthamiana* leaves. BRI1 associated with all tested GRFs, but not with free GFP (Fig. 1B). For further analysis we selected GRF6 to represent all GRFs as this non-epsilon isoform is the closest homolog of the epsilon group (Obergfell et al., 2024). The association between GRF6 and BRI1^K911E^-CD, a kinase-dead BRI1, was dramatically weaker than that with BRI1-CD shown by the GST pull-down assays (Fig. 1C) indicating the necessity of kinase activity for BRI1 and 14-3-3 protein interaction. To further confirm the direct interaction between BRI1 and GRF6, we performed mating-based split-ubiquitin (mbSUS) assay. Yeast cells transformed with *pBT3-STE-BRI1* and *pPR3N-GRF6* grew on SD-TLHA selective medium, whereas those transformed with the negative controls did not (Fig. 1D). Altogether our data demonstrate that GRFs can bind BRI1 directly depending on BRI1 kinase activity.

To further determine whether BRI1 associates more strongly with 14-3-3s in its nascent or active state, we generated transgenic lines overexpressing both *GRF6-HA* and *BRI1-GFP*. Seedlings were first treated with BRZ, followed by treatment with or without the most biologically active BR, brassinolide (BL). We observed an increased association between BRI1 and GRF6 after BL treatment compared with the seedlings treated with BRZ alone (Fig. 1E). In contrast, BL treatment led to increased association between BKI1 and GRF6, but decreased association between BES1 and GRF6 in the transgenic lines overexpressing *GRF6-3xFlag* in *35S::BKI1-GFP*/Col-0 and *GRF6-HA* in *35S::BES1-S-GFP*/Col-0, respectively (Fig. 1E).

### GRF6 binds BRI1 at S858 and T872 in the juxtamembrane domain

Because the BRI1-CD used in the pull-down assays (Fig. 1A) contained a juxtamembrane domain (JM), a kinase domain (KD), and a C-terminal domain (CTD), we sought to investigate which domain is sufficient to mediate BRI1 interaction with GRF6. To that aim, we performed pull-down assays by using different truncations of BRI1 fused to GST and HIS-GRF6. GRF6 was pulled down by BRI1-JK, containing both JM and KD, but not by BRI1-KC, containing both KD and CTD, or BRI1-K, containing only the KD domain, which indicates that the JM of BRI1 mediates a direct interaction with GRF6 (Fig. 2, A and B).

**Figure 2.**
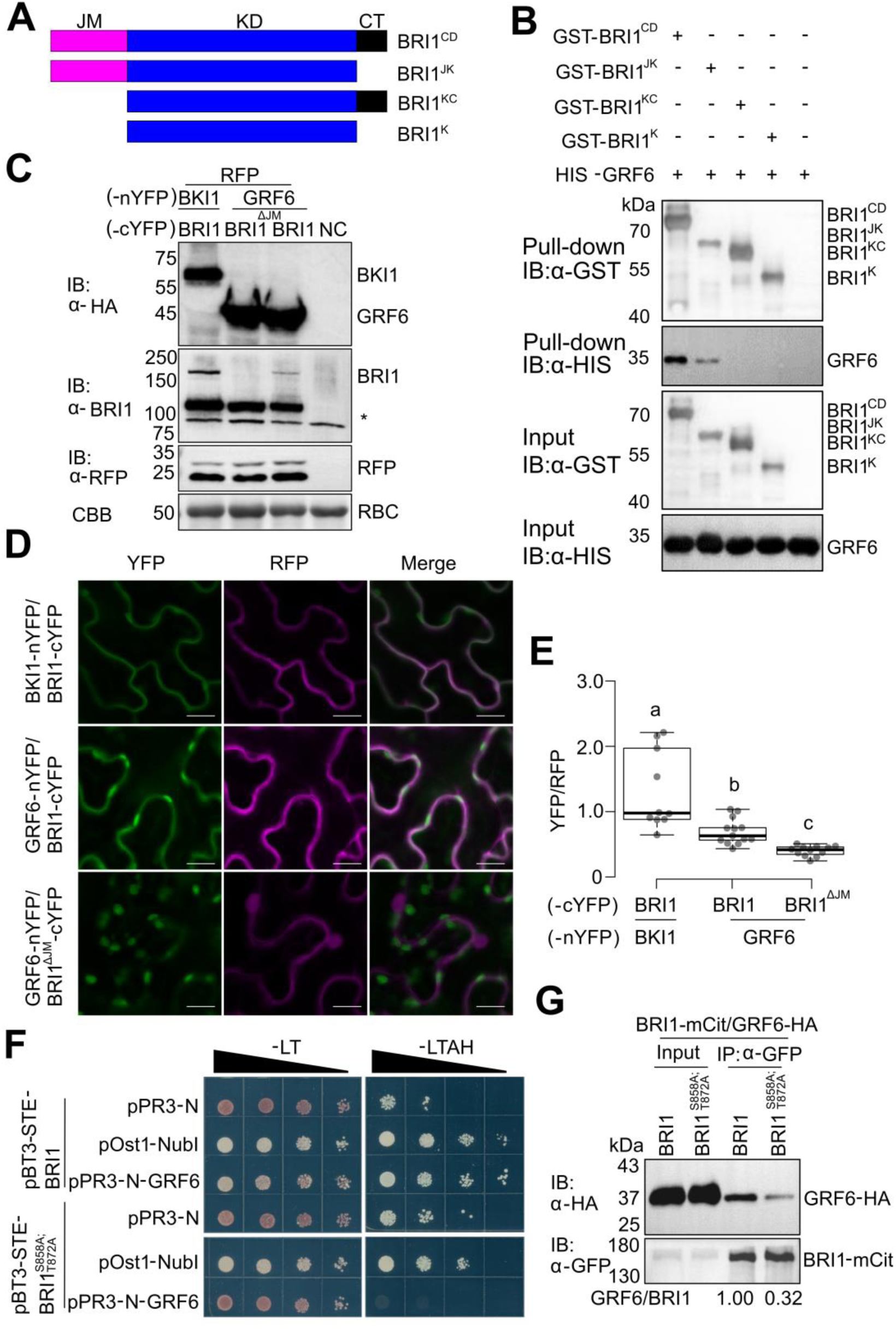
S858 and T872 in the JM domain of BRI1 are required for the 14-3-3-binding to BRI1. **A**) Diagrams showing different truncated forms of BRI1-CD. **B**) GRF6 interaction with BRI1 *in vitro* in the JM domain. HIS-fused GRF6 was incubated with glutathione beads alone or coupled with the GST-fused truncated BRI1-CD, as shown in (**A**). The beads were collected, washed, and subjected to IB with α-GST and α-HIS antibodies. Protein inputs were also determined using α-GST and α-HIS IBs. **C**-**E**) rBiFC analysis of BRI1 and BRI1^ΔJM^ (BRI1 with JM deletion) with GRF6 in *N. benthamiana* leaf epidermal cells. The combination BKI1-nYFP/BRI1-cYFP was used as positive control. The expression levels of each interested protein were subjected to IB using α-HA and α-BRI1 antibodies (**C**). * indicates the specific band originated from degradation of introduced Arabidopsis BRI1. The internal control was blotted with α-RFP antibody. Protein loading was determined by Coomassie brilliant blue (CBB) staining. Quantification of YFP against RFP fluorescence signal ratios for different combinations is shown in (**D** and **E**). *n*=10, *n*, cell number. Scale bar, 10 µm. Different letters indicate significant differences (*P* < 0.05, one-way ANOVA followed by Tukey’s test). **F**) Mating based split-ubiquitin assay of the interactions between BRI1 or BRI1^S858A;T872A^ and GRF6. *pBT3-STE-BRI1/pPR3N* and *pBT3-STE-BRI1^S858A;T872A^/pPR3N* are self-activation controls for BRI1 and BRI1^S858A;T872A^, *pBT3-STE-BRI1/pOst1-NubI* and *pBT3-STE-BRI1^S858A;T872A^/pOst1-NubI* are positive controls for BRI1 and BRI1^S858A;T872A^, and *pBT3-STE-BRI1/pPR3N-GRF6* and *pBT3-STE-BRI1^S858A;T872A^/pPR3N-GRF6* show the interactions between BRI1 and GRF6 or between BRI1^S858A;T872A^ and GRF6. Ten-fold serial dilutions of the saturated cultures were spotted onto plates. **G**) Impaired association of BRI1 with GRF6 in *Arabidopsis* by the S858A and T872A mutations in BRI1. Rosetta leaves of 3-week-old Arabidopsis transgenic lines co-expressing mCitrine (mCit)-tagged BRI1 or BRI1^S858A;T872A^ and HA-tagged GRF6 were harvested for co-IP using α-GFP-coated beads. Proteins were subjected to IBs using α-HA and α-GFP antibodies. The signal ratios of α-HA intensity (GRF6) versus α-GFP intensity (BRI1) are shown below the blots. The blots in **B**), **C**), **D**) and **G**) were performed twice, and similar results were obtained.

To validate the importance of JM for the BRI1-GRF6 interaction, a ratiometric bimolecular fluorescence complementation (rBiFC) assay was carried out in *Nicotiana benthamiana* (Grefen and Blatt, 2012). C-terminally tagged BRI1 (fused with the C-terminal fragment of yellow fluorescent protein, cYFP) combined with C-terminally tagged BKI1 (fused with the N-terminal fragment of YFP, nYFP) was used as a positive control (Liu et al., 2020). C-terminally tagged GRF6 (GRF6-nYFP) was combined with C-terminally tagged BRI1 or BRI1 with JM deletion (BRI1^ΔJM^). With all tested proteins expressed at comparable levels (Figure 2C), BKI1-nYFP/BRI1-cYFP, GRF6-nYFP/BRI1-cYFP, and GRF6-nYFP/BRI1^ΔJM^-cYFP exhibited strong, weak, and no interactions, as indicated by the YFP/RFP signal ratios, respectively (Fig. 2, D and E). Thus, deletion of the JM in BRI1 abolished its interaction with GRF6 *in vivo*, confirming the findings of the GST pull-downs (Fig. 2B). Notably, higher laser power and more open pinhole were used to monitor the association between GRF6 and BRI1^ΔJM^ (Fig. 2C), under which condition chloroplast autofluorescence was also detected.

Most 14-3-3 targets contain motif(s) characterized by the amino acid sequences including (R/K)XX(pS/pT)XP, (R/K)XXX(pS/pT)XP, (pS/pT)X_1-2_-COOH, or YpTV (Johnson et al., 2010; Ormancey et al., 2017; Huang et al., 2022). To identify the residue(s) that mediate(s) BRI1 interaction with GRF6 in JM, we looked for potential 14-3-3 binding motifs. S858, T872, and T880 had previously been identified as phosphorylated residues *in vitro* and *in vivo* (Wang et al., 2005; Wang et al., 2008), of which S858 and T872 and the surrounding residues, VKEAL(p)S858INLAA and PLRKL(p)T872FADLL, match the cargo recognition motif of 14-3-3s. ITC experiments with GRF8 and synthetic BRI1 peptides containing these motifs revealed that GRF8 only binds the phosphorylated RKL(p)T872FA motif, but not the phosphorylated KEAL(p)S858IN motif, nor the non-phosphorylated peptides (Obergfell et al., 2024). Although only (p)T872 was identified as GRF8 recognition site using ITC, because 14-3-3s form dimers to bind to their client protein at two sites, as in the case for BZR1 (Gampala et al., 2007) and BKI1 (Wang et al., 2011), it is still possible that both S858 and T872 are required for 14-3-3s to bind BRI1 *in vivo*.

To confirm the importance of S858 and T872, we first performed mbSUS assay by combining GRF6 with wild-type BRI1 or BRI1 with S858 and T872 mutated to alanine (A), namely BRI1^S858A;T872A^ (Fig. 2F). Yeast cells containing *pBT3-STE-BRI1^S858A;T872A^* and *pPR3-N-GRF6* did not grow on SD-LTAH medium, but in contrast, cells containing *pBT3-STE-BRI1* and *pPR3-N-GRF6* (positive control) were viable. To verify the involvement of S858 and T872 in the BRI1-GRF6 interaction in Arabidopsis, we introduced the C-terminally tagged with mCitrine (mCit) BRI1^S858A;T872A^ mutant driven by native promoter into the *bri1* null mutant, and use *pBRI1:BRI1-mCit/bri1* (Martins et al., 2015) as the control. GRF6-HA was further transformed into homozygous *pBRI1:BRI1-mCit/bri1* and *pBRI1:BRI1^S858A;T872A^-mCit/bri1* lines, which were used for co-IP assays. The association between GRF6 and BRI1 was weakened by the S858A and T872A mutation (Fig. 2G).

As full activation of BRI1 in the presence of BRs relies on its kinase activity and association with BAK1 (Wang et al., 2001; Li et al., 2002; Nam and Li, 2002; Wang et al., 2008), we examined the effect of the S858A and T872A mutations on BRI1 kinase activity and BRI1-BAK1 association. The recombinant BRI1-CD mutant protein (GST-BRI1^S858A;T872A^-CD) was compared with the wild-type BRI1 and the kinase-dead BRI1^K911E^-CD mutant for their ability to autophosphorylate using anti-phosphoserine and threonine (α-pS/T) and anti-phosphotyrosine (α-pY) antibodies (Supplementary Fig. S1A). The BRI1^S858A;T872A^-CD mutant behaved similarly to the wild-type BRI1-CD. Notably, the single BRI1^S858A^-CD migrated faster than the BRI1-CD resembling the kinase inactive BRI1^K911E^-CD mutant (Supplementary Fig. S1B), whereas the T872A mutation was previously shown to increase the kinase activity of BRI1 (Wang et al., 2005). We thus concluded that the two S858 and T872 phosphorylation sites are interdependent and hence they both might be required for the 14-3-3 recognition and binding *in vivo*. This is supported by the comparison of the predicted structures of BRI1-CD (non-phosphorylated form)/GRF6 dimmer and BRI1^S858/T872-P^-CD (S858 and T872 are phosphorylated)/GRF6 dimmer by AlphaFold (Supplemental Figure S1, C and D), as more polar contacts were built between BRI1^S858/T872-P^-CD and GRF6 than between BRI1-CD and GRF6 (Supplemental Figure S1, C and D).

Next, using the GST pull-down assay, we demonstrated that BRI1 and BRI1^S858A;T872A^ bind to BAK1 to almost the same extent (Supplementary Fig. S1E). As a verification in *Arabidopsis*, we performed co-IP assays in the *pBRI1:BRI1-mCit/bri1* and *pBRI1:BRI1^S858A;T872A^-mCit/bri1* lines. To trigger the interaction between BRI1 and BAK1, the seedlings were first treated with BRZ, and then with BL. The associations between BRI1 and BAK1, as well as between BRI1^S858A;T872A^ and BAK1, were comparable (Supplementary Fig. S1F). Taken together, the impaired 14-3-3 binding did not interfere with the BRI1 kinase activity or the BRI1-BAK1 interaction. Accordingly, these data imply that both S858 and T872 phosphorylation sites contribute to GRF6 binding to BRI1.

We next tested whether 14-3-3 binding to BRI1 has any effect on its association with BAK1 and phosphorylation of BAK1. We performed co-IP assays by using *pBRI1:BRI1-mCit/bri1* and *p35S:GRF6-HA*;*pBRI1:BRI1-mCit/bri1* seedlings (Supplementary Fig. S3A). Intriguingly, the presence of GRF6 did not influence the association between BRI1 and BAK1 in the presence of BL (Supplementary Fig. S2A). BRI1 kinase activity was measured by incubating GST-BRI1-CD or HIS-BAK1^K317E^-CD with or without HIS-GRF6. As detected by α-pS/T IB, the signal intensity of phosphorylated BAK1^K317E^-CD in the presence of HIS-GRF6 was similar to that without GRF6 (Supplementary Fig. S2B), indicating that GRF6 binding to BRI1 does not affect BRI1 kinase activity nor its association with BAK1.

### Interference with 14-3-3 binding sites in BRI1 compromised BR signaling

To determine the effect of the S858A and T872A mutations on BR signaling, two homozygous *pBRI1:BRI1^S858A;T872A^-mCit/bri1* lines (lines 3-3 and 7-3) with *BRI1* transcript levels comparable to or higher than that of the complemented *pBRI1:BRI1-mCit/bri1* line (Fig. 3A) were selected for phenotypic analysis. Surprisingly, BRI1^S858A;T872A^ protein levels were reduced when compared to that of the control (Fig. 3B), suggesting that BRI1 protein stability may be impaired by the S858A and T872A mutations. Consistently, *bri1* was only partially complemented by the *pBRI1:BRI1^S858A;T872A^-mCit* construct as shown by the reduced rosette leaf area (Fig. 3C).

**Figure 3.**
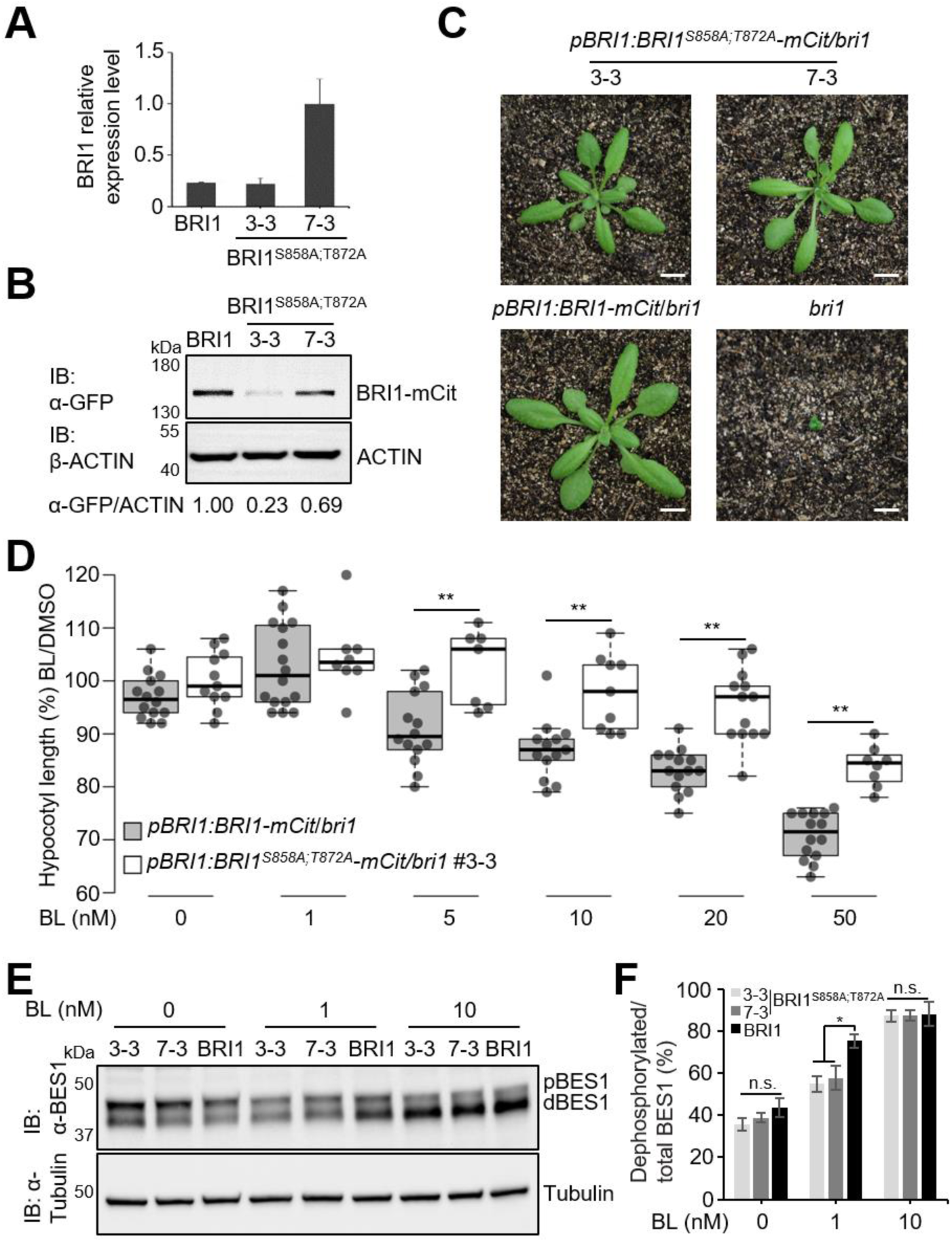
BRI1 carrying S858A and T872A mutations only partially complemented the *bri1* null mutant. **A** and **B**) Molecular characterization of Arabidopsis transgenic lines harboring S858A and T872A mutations in BRI1. **A**) Real-time quantitative reverse transcription-PCR analysis of *BRI1*. Total RNA was isolated from 5-day-old seedlings. *ACTIN2* expression was used as an internal control. Error bars indicate standard deviation (SD); *n* = three biological replicates. **B**) Immunoblot (IB) analysis of BRI1 expression. Protein inputs were equilibrated by IB assay with the β-ACTIN antibody (bottom). **C**) Growth phenotypes of soil-grown 22-day-old homozygous plants. Scale bar, 2 cm. **D**) Sensitivity to brassinolide (BL) of transgenic lines shown by hypocotyl length. Hypocotyl length (normalized to the DMSO control) of 5-day-old seedlings grown in the dark and in the presence of increasing BL concentrations. For each line at each concentration, at least seven seedlings were measured. **E**) Five-day-old seedlings were pre-treated in ½MS for 1 h, followed by treatment with DMSO, 1 nM BL, or 10 nM BL for 0.5 h. Seedlings were further subjected to IB assay using α-BES1 antibody. Protein loading was equilibrated by IB with the α-Tubulin antibody (bottom). The percentage of dephosphorylated BES1 relative to the total BES1 from three biological replicates is shown in (**F**). \**P* < 0.05, \*\**P* < 0.01, Student *t*-test in (**D**) and (**F**).

For quantification of BR sensitivity of the transgenic lines, the length of dark-grown hypocotyls of line 3-3 and the complemented control line in the presence of BL was measured. Line 3-3 was less sensitive to BL at concentrations of 5, 10, 20, and 50 nM than the control (Fig. 3D). Dephosphorylation of BES1 by exogenous BL is commonly used as a BR signaling activation readout (Yin et al., 2002). In agreement with the hypocotyl growth assay results, application of BL at 1 nM for 0.5 h, dephosphorylated BES1 (dBES1) to a lesser extent in both lines 3-3 and 7-3 than in the control (Fig. 3, E and F). Collectively, the *pBRI1:BRI1^S858A;T872A^-mCit/bri1* lines were less sensitive to exogenous BRs probably due to the reduced PM levels of BRI1 (Fig. 3, D-F).

### Depletion of 14-3-3 binding in BRI1 led to BRI1 polyubiquitination

BRI1 undergoes polyubiquitination directly mediated by PUB12 and PUB13 (Zhou et al., 2018) and is further degraded in the vacuole (Martins et al., 2015). The proteasome inhibitor MG132 may stabilize BRI1, as BRI1-PUB13 association was more obvious in the presence of MG132 (Zhou et al., 2018). MG132 also stabilized BRI1^S858A;T872A^ as that for BRI1 in the complemented line (Fig. 4A), reflecting, to some extent, the transcript levels of *BRI1^S858A;T872A^*or *BRI1* in each line (Fig. 3A). This indicates that the impaired protein stability for BRI1^S858A;T872A^ may be a result of stronger ubiquitination modification than the wild-type BRI1.

**Figure 4.**
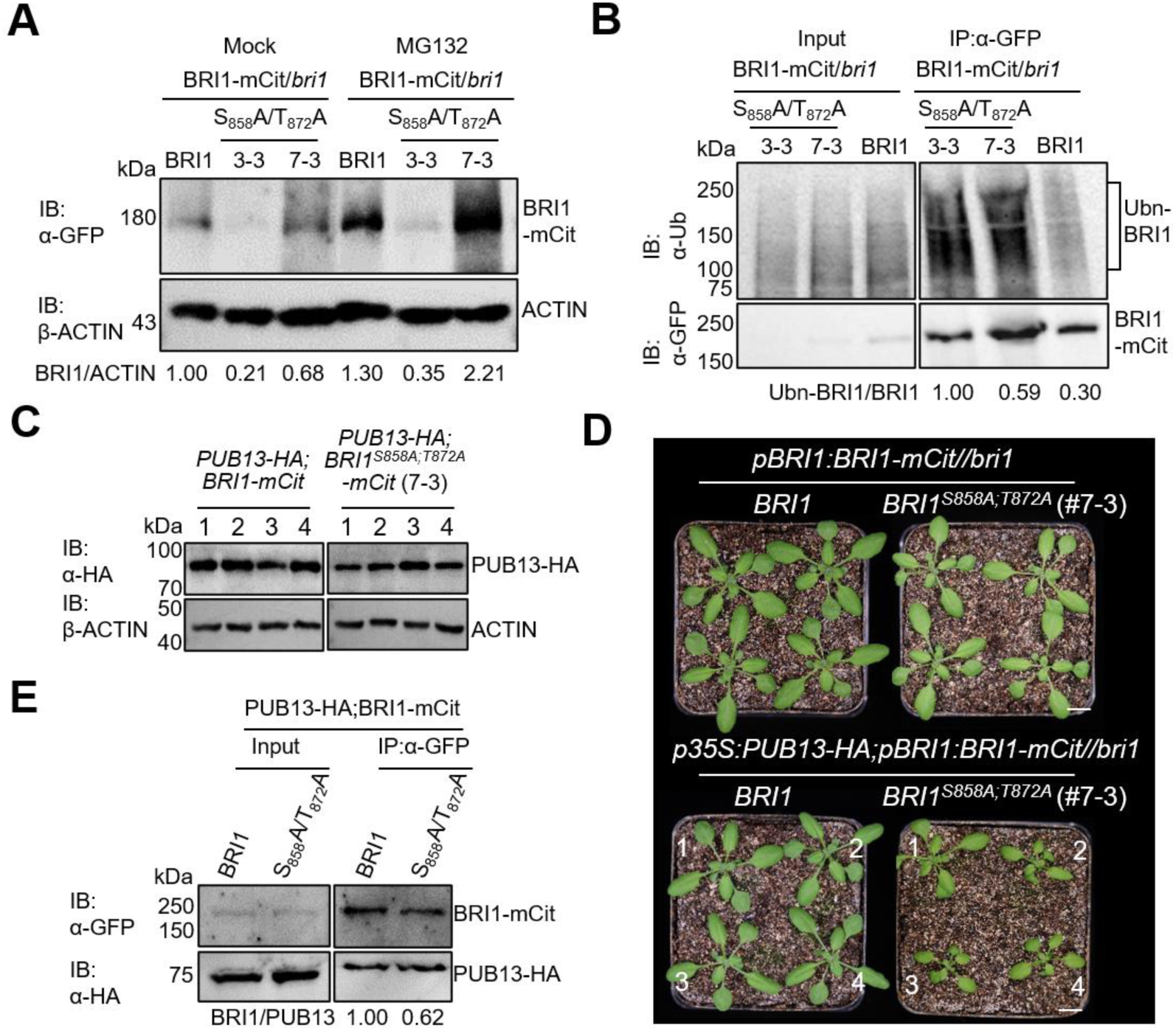
S858A and T872A mutations reduced the protein abundance of BRI1. **A**) BRI1^S858A;T872A^ abundance depends on the 26S proteasome. Six-day-old seedlings of the indicated lines were treated with 50 µM MG132 or an equal volume of DMSO for 5 h before harvesting for total protein isolation. BRI1 protein amount was determined by IB with the α-BRI1 antibody (top), and protein inputs were equilibrated by IB with the β-ACTIN antibody (bottom). The signal ratios of BRI1 versus ACTIN are shown below the blots. **B**) Increased BRI1 ubiquitination in BRI1^S858A;T872A^. Immunoprecipitation (IP) was performed using α-GFP-coated magnetic beads on solubilized microsomal fraction proteins isolated from 6-day-old seedlings of the indicated transgenic lines after treatment with MG-132 (50 µM) for 5 h and subjected to IB assay with α-ubiquitin (top) and α-GFP (bottom). The ratios between ubiquitinated BRI1 and total BRI1 (Ub_n_-BRI1/BRI1-mCit) were calculated and are shown below the blots. **C**) IB analysis of PUB13-HA expression. Total proteins isolated from the indicated T1 transgenic plants were detected using an α-HA antibody (top). Protein inputs were equilibrated by immunoblot (IB) assay with the β-ACTIN antibody (bottom) **D**) Growth phenotypes of the soil-grown 23-day-old T1 *p35S:PUB13-HA;pBRI1:BRI1-mCit/bri1* and *p35S:PUB13-HA;pBRI1:BRI1^S858A;T872A^-mCit/bri1* (#7-3) plants, together with the indicated control lines. Scale bar, 1 cm. **E**) Enhanced association between BRI1 and PUB13 in *Arabidopsis* by the S858A and T872A mutations in BRI1. 10-day-old Arabidopsis seedlings co-expressing mCit-tagged BRI1 or BRI1^S858A;T872A^ and HA-tagged PUB13 were harvested for co-IP using α-GFP-coated beads. Proteins were subjected to IB assays using α-HA and α-GFP antibodies. The signal ratios of α-GFP intensity (BRI1) versus α-HA intensity (PUB13) are shown below the blots. The blots in **A**), **B**) and **E**) were performed twice, and similar results were obtained.

To determine BRI1^S858A;T872A^ ubiquitination level, we immunoprecipitated BRI1 (or BRI1^S858A;T872A^)-mCit from the three lines by using the isolated microsomal fractions and probed them with α-ubiquitin antibodies (Fig. 4B). An increased molecular mass smear of approximately 100–250 kDa was detected in all samples, corresponding to polyubiquitinated BRI1. Corresponding with the decreased BRI1 stability for BRI1^S858A;T872A^ and measured by the relative signal intensity ratio between ubiquitinated BRI1 (Ubn-BRI1) and immunoprecipitated BRI1, the BRI1 ubiquitination intensity was stronger in the two *pBRI1:BRI1^S858A;T872A^-mCit/bri1* transgenic lines than in the control. The ratio between the ubiquitination of BRI1 and the total BRI1 protein (Ub_n_-BRI1/BRI1-mCit) was 2-fold and 1-fold higher in lines 3-3 and 7-3, respectively (Fig. 4B).

BRI1 polyubiquitination is mediated by PUB12 and PUB13. To investigate the connection between 14-3-3 binding to BRI1 and PUB13-mediated BRI1 ubiquitination, we overexpressed PUB13-HA in *pBRI1:BRI1-mCit;bri1* and *pBRI1:BRI1^S858A;T872A^-mCit*/*bri1* (#7-3) (Fig. 4C). Compared with *p35S:PUB13-HA/pBRI1:BRI1-mCit;bri1* plants, most *p35S:PUB13-HA/pBRI1:BRI1^S858A;T872A^-mCit;bri1* (#7-3) plants exhibited a *bri1* mutant-like phenotype (Fig. 4D), suggesting that BRI1^S858A;T872A^ is hypersensitive to *PUB13* overexpression, and that BRI1^S858A;T872A^ may be degraded more than the wild-type BRI1 in a PUB13-dependent manner. The hypersensitivity of BRI1^S858A;T872A^ to *PUB13* overexpression correlated with the increased association between BRI1^S858A;T872A^ and PUB13 than that between BRI1 and PUB13 as shown by the co-IP assays (Figure 4E).

### Absence of 14-3-3 binding destabilized BRI1 in the plasma membrane

As ubiquitination of BRI1 regulates its endocytosis (Martins et al., 2015; Zhou et al., 2018) and the S858A and T872A mutations cause increased BRI1 ubiquitination (Fig. 4B), we examined whether the PM and vacuole targeting of BRI1^S858A;T872A^ were altered. To this end and because the fluorescence signals were too weak to be detected in the *pBRI1:BRI1^S858A;T872A^-mCit/bri1* and *pBRI1:BRI1-mCit/bri1* lines, we generated overexpression lines expressing BRI1-GFP and BRI1^S858A;T872A^-GFP from the *35S* promoter. We evaluated the expression level of BRI1 (or BRI1^S858A;T872A^)-GFP and selected two lines for each construct with comparable expression levels for further characterization (Supplementary Fig. S3A). Both *p35S:BRI1-GFP*/Col-0 and *p35S:BRI1^S858A;T872A^-GFP*/Col-0 lines display phenotypes of enhanced BR signaling (Supplementary Fig. S3B). But no obvious differences were observed between *p35S:BRI1-GFP*/Col-0 and *p35S:BRI1^S858A;T872A^-GFP*/Col-0 lines (Supplementary Fig. S3B).

Next, to determine whether the mutation would increase BRI1 internalization, we utilized several chemical treatments. Using cycloheximide (CHX) to inhibit *de novo* protein synthesis, we assessed the PM pool of BRI1 in the root meristem of 5-day-old seedlings. The ratio of PM to intracellular fluorescence intensity in *p35S:BRI1^S858A;T872A^-GFP*/Col-0 plants was significantly lower than that in *p35S:BRI1-GFP*/Col-0 seedlings (Fig. 5, A and B), suggesting an increase in intracellular BRI1^S858A;T872A^. To decipher further the contribution of each of the two residues to BRI1 internalization, we generated *p35S:BRI1^S858A^-GFP*/Col-0 and *p35S:BRI1^T872A^-GFP*/Col-0 lines. Endocytosis of BRI1^S858A^ and BRI1^T872A^ is impaired and enhanced, respectively (Supplementary Fig. S4), which may be largely determined by their decreased and increased kinase activity (Supplementary Fig. S1B) (Wang et al., 2005).

**Figure 5.**
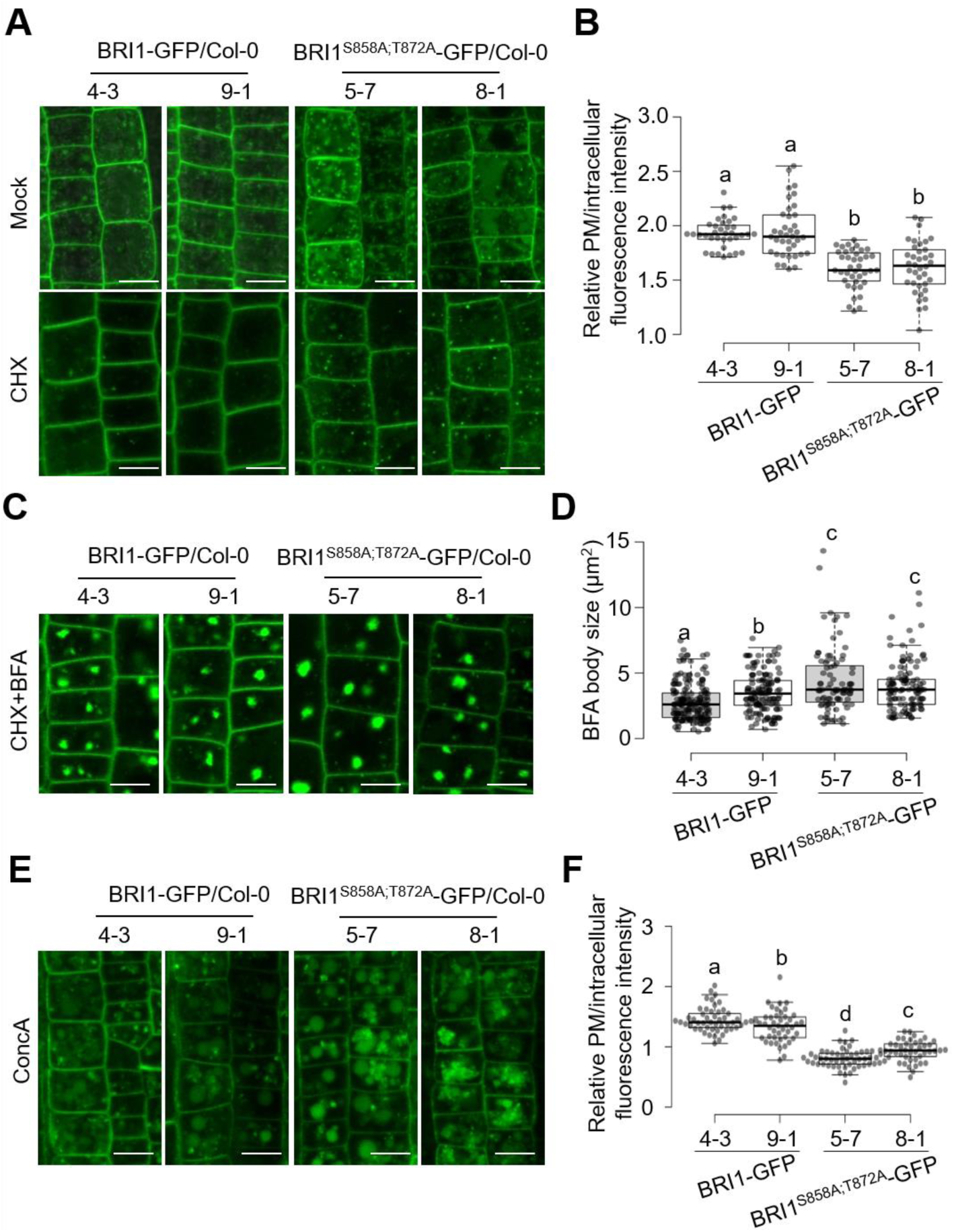
S858A and T872A mutations in BRI1 decreased the plasma membrane (PM) pool of BRI1 and increased its internalization, and targeting to the vacuole. **A** and **C**) Images of epidermal cells from root meristems of 5-day-old transgenic Arabidopsis seedlings expressing WT or S858A;T872A-mutated BRI1 tagged with GFP pretreated with cycloheximide (CHX) (50 µM) for 1.5 h (**A**) or pretreated with CHX (50 µM) for 1 h, followed by a combined treatment with CHX (50 µM) and BFA (50 µM) for 30 min (**C**). **B** and **D**) Measurements of the relative PM versus intracellular BRI1-GFP fluorescence intensity and BFA body size derived from the images displayed in (**A**) and (**C**). At least 36 cells from five seedlings were measured for each transgenic line. **E**) Sensitivity to ConcA of BRI1-GFP/Col-0 and BRI1^S858A;T872A^-GFP/Col-0 plants. Five-day-old seedlings of the indicated lines were exposed to ConcA (1 µM) for 2 h. **F**) Measurements of the relative PM versus intracellular BRI1-GFP fluorescence intensity from the images displayed in (**E**). Similar confocal detection settings were used to compare different lines under different chemical treatments. Scale bar, 10 µm in (**A**), (**C**), and (**E**). Different letters in (**B**), (**D**), and (**F**) indicate significant differences (*P* < 0.05, one-way ANOVA followed by Tukey’s test).

By passing through the trans-Golgi network/early endosomes (TGN/EEs), internalized BRI1 is targeted to vacuoles for degradation or recycled back to the PM (Irani et al., 2012). To better visualize the internalized BRI1, we took advantage of another chemical Brefeldin A (BFA) to inhibit endosomal trafficking, which leads to the accumulation of PM proteins in BFA bodies (Geldner et al., 2001). Consistent with the increased intracellular BRI1^S858A;T872A^ (Fig. 5, A and B), the GFP signal in BFA bodies was stronger in the BRI1^S858A;T872A^-GFP than that in BRI1-GFP seedlings, as indicated by larger BFA bodies (Fig. 5, C and D).

To prevent BRI1 degradation in vacuoles (Martins et al., 2015), we applied Concanamycin A (ConcA), a vacuolar ATPase inhibitor, which allows visualization of accumulated BRI1-GFP or BRI1^S858A;T872A^-GFP in vacuoles. ConcA treatment induced a dramatic accumulation of BRI1^S858A;T872A^-GFP in the vacuole compared to BRI1-GFP (Fig. 5E), thus revealing an enhanced vacuolar delivery of BRI1^S858A;T872A^ that may be triggered by ubiquitination modification. In addition to the GFP signal enhancement in vacuoles, larger ConcA bodies were present in *p35S:BRI1^S858A;T872A^-GFP*/Col-0 than in *p35S:BRI1-GFP*/Col-0 seedlings (Fig. 5E).

### Loss-of-function or silencing 14-3-3s reduced BRI1 abundance in the plasma membrane

To gain further insight into the function of 14-3-3s in BR signaling we used a quadruple *grf1,4,6,8* mutant also known as *klpc* (van Kleeff et al., 2014) (Supplementary Fig. S6A), in which Kappa (κ; *GRF8*), Lambda (λ; *GRF6*), Phi (φ; GRF4), and Chi (χ; GRF1) non-epsilon isoforms are mutated, hereinafter referred to as *grf1,4,6,8* (Supplementary Fig. 5A). The abundance of BRI1 in *grf1,4,6,8* was not altered (Fig. 6A). Considering the possible functional redundancy of 14-3-3s, and the ubiquitous expression of the non-epsilon isoforms suggesting low specificity (Keicher et al., 2017), we applied the CRISPR-Cas9 genome editing strategy to generate higher-order mutants in the *grf1,4,6,8* background. To reduce the risk of low editing efficiency, we designed two gRNAs for each of the remaining four non-epsilon GRFs, *GRF2*, *GRF3*, *GRF5,* and *GRF7*, and introduced the genome-editing cassette into *grf1,4,6,8*. From the mutant library, we succeeded in obtaining an octuple *grf1,2,3,4,5,6,7,8* mutant, creating pre-mature terminations in *GRF2*, *GRF5*, and *GRF7* and a 6-bp deletion in *GRF6* leading to 2 amino acid deletion (Supplementary Fig. 5B). In contrast to *grf1,4,6,8*, the protein amount of BRI1 was clearly reduced in *grf1,2,3,4,5,6,7,8* (Fig. 6A), which was not a result of reduced *BRI1* transcription (Supplementary Fig. 5C). In agreement, the soil-grown *grf1,2,3,4,5,6,7,8* plants were smaller than both WT and *grf1,4,6,8* mutant (Supplementary Fig. 5D), phenocopying a weak *bri1* mutant (Sun et al., 2017).

**Figure 6.**
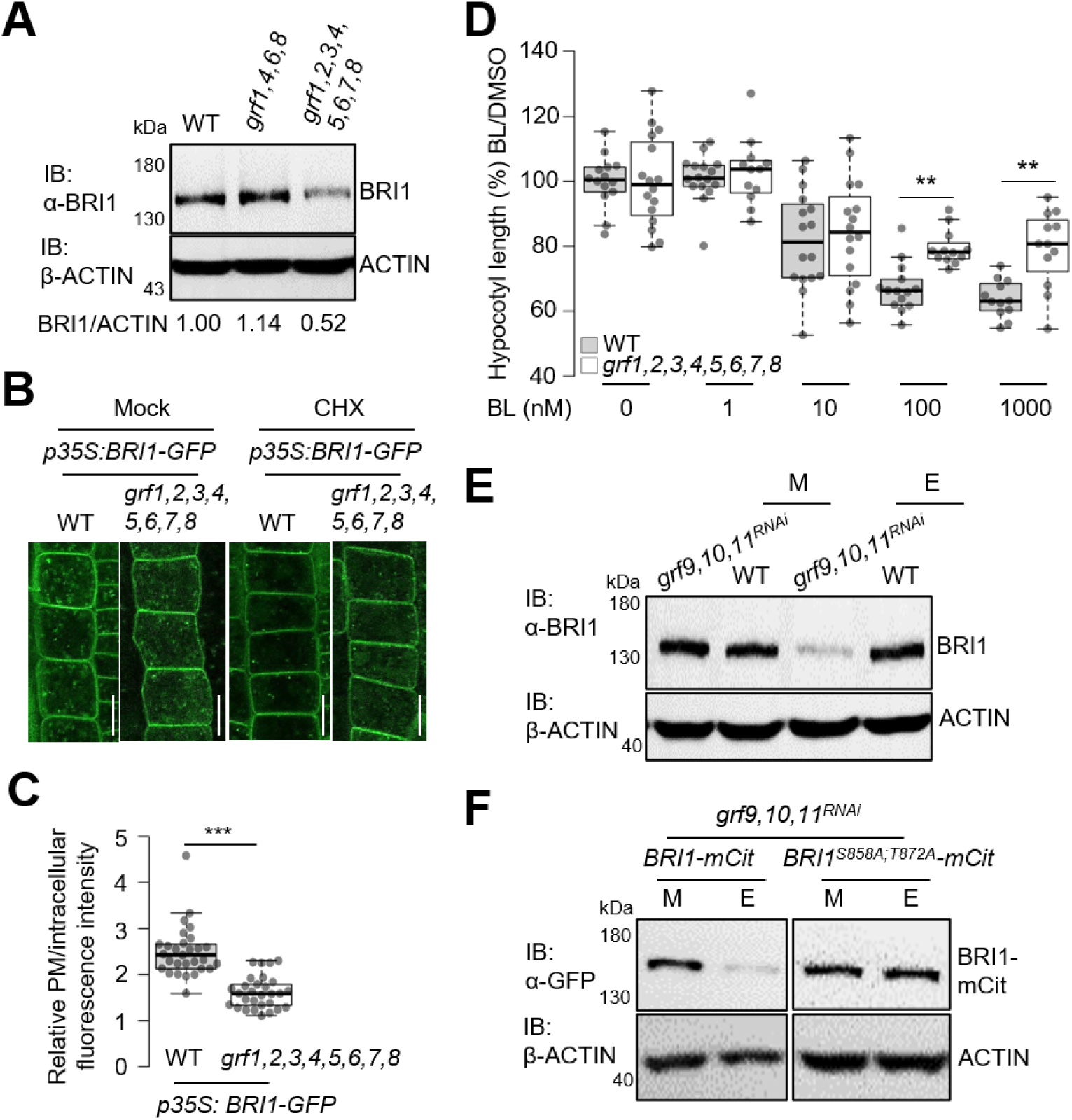
Knocking out *GRF2*, *GRF3*, *GRF5*, and *GRF7* in *grf1,4,6,8* and silencing *GRF9*, *GRF10,* and *GRF11* (*grf9,10,11^RNAi^)* resulted in BRI1 degradation. **A**) Reduced BRI1 stability in *grf1,2,3,4,5,6,7,8* compared to the controls. Total proteins were isolated from 6-day-old seedlings of the indicated lines. The amount of BRI1 protein was determined by IB assay with the α-BRI1 antibody (top), and IB with the β-ACTIN was used as loading control (bottom). The signal ratios of BRI1 versus ACTIN are shown below the blots. **B** and **C**) Enhanced BRI1 endocytosis in *grf1,2,3,4,5,6,7,8* than in WT background. **B**) Images of epidermal cells from root meristems of 6-day-old screened T1 Arabidopsis seedlings expressing BRI1 tagged with GFP pretreated with cycloheximide (CHX) (50 µM) for 1.5 h. **C**) Measurements of the relative PM versus intracellular BRI1-GFP fluorescence intensity derived from the images displayed in the right panel in (**B**). At least 30 cells from four seedlings were measured for each background. **D**) Sensitivity to BL of transgenic lines shown by hypocotyl length. Hypocotyl length (normalized to the DMSO control) of 5-day-old seedlings grown in the dark and in the presence of increasing BL concentrations. For each line at each concentration, at least 12 seedlings were measured. \*\**P* < 0.01, Student *t*-test. **E** and **F**) Immunoblotting (IB) analysis of BRI1 and BRI1-mCitrine (mCit) abundance. Total protein was isolated from 7-day-old seedlings of the indicated lines grown in a medium containing 0.1% (v/v) ethanol (E) or mock (M). The total protein amount isolated from homozygous *bri1* was used as negative control to demonstrate the specificity of the α-BRI1 antibody. BRI1 and BRI1-mCit were detected using α-BRI1 and α-GFP antibodies, respectively (top). ACTIN was used as loading control by IB with the β-ACTIN antibody (bottom). Scale bar in (**B**), 10 µm.

To characterize BRI1 internalization behavior, we overexpressed *p35S:BRI1-GFP* in *grf1,2,3,4,5,6,7,8*. We further assessed the PM pool of BRI1 in the root meristem of 6-day-old seedlings after CHX treatment. The ratio of PM to intracellular fluorescence intensity in *BRI1-GFP/grf1,2,3,4,5,6,7,8* seedlings was lower than that in *p35S:BRI1-GFP*/Col-0 seedlings (Fig. 6, B and C), indicating an increase in BRI1 internalization in *BRI1-GFP/grf1,2,3,4,5,6,7,8*, and the reason for impaired BRI1 protein stability in *grf1,2,3,4,5,6,7,8*. In line with the reduced PM pool of BRI1, *grf1,2,3,4,5,6,7,8* seedlings were less sensitive than the WT seedlings to only the higher concentrations (100 and 1000 nM) of BL, as determined by measuring the hypocotyl length of the dark-grown seedlings treated with BL (Fig. 6, D).

Given that the epsilon isoforms Mu (μ; GRF9), Epsilon (ε; GRF10), and Omicron (ο; GRF11) can also bind BRI1 (Fig. 1, A and B), and that they are also ubiquitously expressed (Keicher et al., 2017), we wondered whether they might also affect BRI1 stability in the same manner as the non-epsilon isoforms. To verify this hypothesis, we used the available ethanol-inducible amiRNA line (Keicher et al., 2017), in which the 14-3-3 isoforms *GRF9*, *GRF10* and *GRF11* are knocked down by an ethanol treatment, hereinafter referred to as *grf9,10,11^RNAi^*(Supplementary Fig. S6A). The transcript level of *BRI1* remained unaffected in *grf9,10,11^RNAi^* similar to that in *grf1,2,3,4,5,6,7,8* (Supplementary Fig. S6B).

Also resembled that in *grf1,2,3,4,5,6,7,8*, BRI1 protein level was lower in ethanol-treated *grf9,10,11^RNAi^* than in the mock, whereas those in wild-type remained unchanged (Fig. 6E). To verify that BRI1 protein stability relies on its association with GRF9, GRF10, and GRF11, we transformed *pBRI1:BRI1-mCit* and *pBRI1:BRI1^S85A8;T872A^-mCit* into *grf9,10,11^RNAi^*. Lines with comparable BRI1-mCit and BRI1^S85A8;T872A^-mCit were selected for further analysis. Similar to the endogenous BRI1 in *grf9,10,11^RNAi^*, when *pBRI1:BRI1-mCit/grf9,10,11^RNAi^* was treated with ethanol, BRI1-mCit protein level decreased (Fig. 6F). Interestingly, the BRI1^S858A;T872A^-mCit levels were insensitive to the ethanol treatment indicating that BRI1 stability governing mediated by GRF9,10, and 11 is through their binding at S858 and T872. In line with the decreased BRI1 protein level, soil-grown ethanol-treated *grf9,10,11^RNAi^* plants were smaller than wild-type plants, phenocopying a weak *bri1* mutant (Supplementary Fig. S6C).

## Discussion

Previous studies on the functions of 14-3-3s in BR signaling have mainly focused on BES1, BZR1 and BKI1. In this study, by mutating the 14-3-3 binding sites in BRI1, and using octuple mutant *grf1,2,3,4,5,6,7,8* and *grf9,10,11^RNAi^*, we demonstrated that both non-epsilon and epsilon 14-3-3 isoforms bind and stabilize BRI1 by antagonizing the ubiquitination modification of BRI1. This, in turn, affected BRI1 ubiquitination and endocytosis. Moreover, a similar scenario of 14-3-3s regulating BRI1 and BR signaling may also exist for other RKs and RK-mediated signaling pathways. Thus, our findings may provide a new perspective on the modulation of RK signaling.

### 14-3-3s-BRI1-BAK1 complex formation, BRI1 ubiquitination and endocytosis

BRI1 heterodimerization with its co-receptor and the subsequent reciprocal transphosphorylation is the prerequisite for BR signal transduction, which requires their spacial proximity. Our *in vitro* and *in vivo* data all suggest that 14-3-3s do not sterically hinder BRI1 and BAK1 interaction. This speculation is also supported by the predicted structures of BRI1-CD/BAK1-CD/GRF6 trimer by AlphaFold, as compared with BRI1-CD/BAK1-CD dimmer, the BRI1-BAK1 interface is only mildly changed in BRI1-CD/BAK1-CD/GRF6 trimer (Supplemental Figure S7). This could be explained by the fact that 14-3-3s bind to BRI1 at the juxta-membrane domain, but not the kinase domain. Since the 14-3-3s form dimers to further bind to their targets, these dimers may target the BRI1-BAK1 heterodimer in the presence of BRs. This hypothesis is supported by evidence showing that 14-3-3s can directly bind to the BAK1 homolog SERK1 (Rienties et al., 2005) and possibly interact with BAK1 in Arabidopsis (Chang et al., 2009). Several putative 14-3-3 binding motifs in the cytoplasmic domain of BAK1 and SERK1 were predicted, including BAK1^323-327^, BAK1^600-610^, SERK1^289-293^, SERK1^412-417^, and SERK1^566-576^, using the eukaryotic linear motif (ELM) server (ELM - Search the ELM resource) (Puntervoll et al., 2003). But, the phosphorylation modification of S_325_ and S_604_ in BAK1, and S_291_, S_415_, and S_570_ in SERK1, have not been reported yet. Thus, non-canonical 14-3-3 binding motifs may exist in BAK1 and SERK1.

BRI1 auto-phosphorylation is required for BRI1-BAK1 association and its full activation. Full activation of BRI1 enhances its interaction with 14-3-3 proteins shown in this study. In its fully activated state, BRI1 interacts with and phosphorylates the E3 ubiquitin ligases PUB12 and PUB13, which in turn ubiquitinate BRI1 for degradation, thus reducing BRI1 abundance at the PM (Zhou et al., 2018). Kinase activity is necessary for BRI1 interaction with 14-3-3s, because the targeted residues of 14-3-3s must be phosphorylated (Obergfell et al., 2024). Binding of 14-3-3s to BRI1 may antagonize PUB12 and PUB13-mediated ubiquitination of BRI1. BRI1 is K63 polyubiquitinated *in vivo,* and ubiquitination promotes its internalization and sorting for vacuolar degradation (Martins et al., 2015; Zhou et al., 2018). One of the major ubiquitination sites in BRI1 is K866, adjacent to S858 and T872, the two 14-3-3s binding sites in BRI1, thus BRI1 polyubiquitination may be sterically hindered by 14-3-3 binding, and vice versa. It may explain the hypersensitivity of *pBRI1:BRI1^S858A;T872A^-mCit/bri1* to PUB13 overexpression shown in our study. Mutation of 14-3-3 binding sites in BRI1 enhanced its ubiquitination and internalization. The ubiquitination of BRI1 is well-known as a prerequisite for its ubiquitin-dependent endocytosis (Martins et al., 2015; Zhou et al., 2018). In addition, BRI1 undergoes AP-2- and TPC-dependent clathrin-mediated endocytosis (Di Rubbo et al., 2013; Gadeyne et al., 2014; Liu et al., 2020) as well as clathrin-independent endocytosis (Wang et al., 2015). Studies with the recently generated tool BRI1^Q^, a BR binding-deficient mutant (Claus et al., 2023), further complicated our understanding of the endocytic machinery mediating BRI1 endocytosis. BRI1 endocytosis was enhanced in *grf1,2,3,4,5,6,7,8* mimicking the phenotype of BRI1^S858A;T872A^ and the artificially ubiquitinated BRI1 (Martins et al., 2015). Regulation of endocytosis by 14-3-3s is not unique for BRI1, because endocytosis of PINOID2 was also enhanced in the *grf9,10,*11^RNAi^ line (Keicher et al., 2017).

### Positive or negative regulators of 14-3-3s in BR signaling depends on where they act

BR signaling is largely determined by the amount of functional BRI1 protein in the PM. Our study revealed that 14-3-3s stabilize BRI1 at PM. However, whether 14-3-3s are general positive or negative regulators of BR signaling remains controversial. As, first, hypersensitivity to BRs resulting from *BRI1* overexpression was reduced in the *14-3-3* triple mutant *grf3,4,10* (Lee et al., 2020), indicating a positive role of 14-3-3s in BR signaling, and second, three quadruple 14-3-3 mutants *grf1,4,6,8*, *grf5,6,7,8*, and *grf1,4,5,7* were weakly less sensitive to BRZ, suggesting that 14-3-3s are negative regulators of BR signaling (Obergfell et al., 2024). Noteworthy, BRZ sensitivity may not be a very clear readout of BR responses as it also affects BR biosynthesis, and BRZ insensitivity is also found in signaling deficient mutants including BR perception mutant *bri1-701*, SERK triple-null mutant *serk1-8 bak1-4 bkk1-1*, and BR biosynthetic null mutant *cpd* (Du et al., 2012). It suggests that reduced BRZ sensitivity does not always equals to more BR signaling. These conflicting findings may be due to isoform binding specificity to a certain extent, for different targets including BRI1, BKI1, BES1, BZR1, and maybe SERKs and BIN2, although GRF8 was shown to bind the phosphorylated short linear motifs in BRI1, BKI1, and BES1 similarly (Obergfell et al., 2024). The conflicting findings may mainly attribute to where 14-3-3s target to, as binding to BRI1, BKI1, or BES1, BZR1 resulting elevated or declined BR signaling. Almost all studies about 14-3-3s are from the non-epsilon isoforms, little was known about how epsilon group 14-3-3s contribute to BR signaling. In addition, their binding efficiency may differ between different 14-3-3 isoforms in terms of the same target.

Our GST pull-down and co-IP assays showed that most of 14-3-3 isoforms could bind to BRI1. BRI1 protein levels were less abundant in both *grf1,2,3,4,5,6,7,8* and *grf9,10,11^RNAi^*. These findings point to a potential additive effect between the non-epsilon and the epsilon 14-3-3 isoform for their recognition and stabilization of BRI1. This phenomenon may not be true for all 14-3-3 targets, as the early flowering phenotype shown in the Flag-tagged BRI1 overexpressed plants in wild type background, lost in the triple *14-3-3* mutant *grf3,4,10*, could not be rescued by introducing any of the three isoforms (Lee et al., 2020). The same may exist for 14-3-3s isoforms binding BKI1, BES1, BZR1, SERKs, BIN2, and maybe more components in BR signaling.

Taken together, drawing a conclusion about whether14-3-3s are generally positive or negative regulators in BR signaling would be unnecessary and unrealistic, as it really depends on where 14-3-3 proteins target to.

### 14-3-3s contribute to RK-mediated signaling and their function may also rely on RKs

14-3-3s target to a large number of proteins in a variety of biological activities (Huang et al., 2021). RKs, as being phosphorylated are also potential targets of 14-3-3s. Large-scale and high throughput interactome assays using each of the 14-3-3 isoforms will map out all the 14-3-3 and RK interaction. Starting from there, the biological functions of these interactions will be of interesting to investigate further.

14-3-3s themselves can be modified by phosphorylation, potentially affecting their sub-cellular localization and client binding (Liu et al., 2017; Aitken, 2011; Gao et al., 2022). BRI1 and BAK1 phosphorylate 14-3-3s in a BR-dependent manner (Chae et al., 2016), and the phosphorylation sites in GRF2 *in planta* has been revealed (Wu et al., 2011). Thus, there may be a feed-back forward effect between the association of 14-3-3s and BRI1 and BAK1 positively regulating BR signaling. Phosphorylated 14-3-3s may also target to other proteins thus participate in other signaling pathways. The crosstalk between BR signaling and others may partially rely on BRI1- and BAK1-mediated 14-3-3 phosphorylation.

## Materials and methods

### Plant materials and growth condition

Arabidopsis *thaliana* L. Heynh., accession Columbia 0 (Col-0) was used as the wild-type. *pBRI1:BRI1-mCit* expressed in *bri1* (*pBRI1:BRI1-mCit/bri1*), *bri1* null mutant (GABI_134E10), *grf1,4,6,8* and *grf9,10,11^RNAi^* were described previously (Jaillais et al., 2011; Martins et al., 2015; Zhou et al., 2018); (van Kleeff et al., 2014; Keicher et al., 2017).

For phenotypic analysis, plants were grown in soil for 22-28 days at 22°C, 50-60% relative humidity, and a 16-h light/8-h dark regime, under a standard daytime light intensity (90 mmol/m^2^/s) from full-spectrum fluorescent light bulbs. For growth on plates or in liquid medium, Arabidopsis seeds were sterilized with sodium hypochlorite and Triton X-100 and sown on plates with half-strength Murashige and Skoog (½MS) medium containing 0.5% (w/v) sucrose, 1% (w/v) agar, and 0.05% (w/v) MES, pH 5.7. After vernalization for 2 days at 4°C, the plates were transferred to the growth chamber at 22°C under a 16-h/8-h light/dark cycle or in the dark after 2 h of light for different periods of time, depending on the experiments. Plants were grown on plates for 6 days for the microsomal protein preparation; for 5 days for the BRI1 internalization assay, transcript analyses, protein abundance analysis, and dark-grown BR and ConcA sensitivity assays; for 4 h followed by 6 more days of treatment with different BL or ConcA concentrations. For Co-IP assays with seedlings, plants were grown on plates for 3 days, then transferred to ½MS liquid medium for 5 days, followed by treatment with BRZ (2 µM) for 3 days and BL (1 µM) for 90 min.

### Molecular cloning

cDNA or genomic DNA from 5-day-old Col-0 seedlings or indicated vectors were used as templates for PCR cloning using the indicated primers listed in Supplementary Table S1. For transient expression in *Nicotiana benthamiana*, *pDONR221-BRI1* was recombined with the *35S* promoter and HA tag into the *pK7m34GW* vector (Karimi et al., 2007) to generate the *pK7m43GW-p35S:BRI1-HA*. The cDNA of *BRI1* was in-fusion cloned into *pCB302-p35S:HA* digested with *Bam*HI and *Stu*I to generate *pCB302-p35S:BRI1-HA*. The gDNA fragment of *GRF6* was cloned into *pDONR221* to generate *pDONR221-GRF6*, which was recombined with the *35S* promoter and *GFP* tag into the *pK7FWG2* to generate *pK7FWG2-p35S:GRF6-GFP*. The truncated version of BRI1 was built by amplification of BRI1^ΔJM^, the fragments comprising extracellular domain and transmembrane domain (ECD+TM), and the kinase domain and C-terminal domain (KD+CT) and cloned into *pUC19* with the LightNing™ DNA Assembly Mix Plus (iScience) to create *pUC19-BRI1^ΔJM^*. The cDNAs encoding *BKI1* or *GRF6* were cloned into *pDONR221-P3P2*, and *BRI1* (or *BRI1^ΔJM^*) into *pDONR221-P1P4*, which were further cloned into *pBiFC-2in1-CC* to generate *pBiFC-BKI1-nYFP-BRI1-cYFP*, *pBiFC-GRF6-nYFP-BRI1-cYFP*, and *pBiFC-GRF6-nYFP-BRI1^ΔJM^-cYFP*. The gDNA fragments of each of the 9 *GRFs* selected including *GRF2*, *GRF3*, *GRF5*, *GRF6*, *GRF9*, *GRF10*, and *GRF11*, were in-fusion cloned into *pCB302-p35S::GFP* digested with *Bam*HI and *Stu*I individually to generate the *pCB302-35S::GRF-GFP* constructs, individually.

For stable expression in Arabidopsis, *pDONR221-BRI1^S858A;T872A^*was obtained by site-directed mutagenesis with *pDONR221-BRI1* (Jaillais et al., 2011) as template. *pDONR221-BRI1^S858A;T872A^*, *pDONRP4P1r-pBRI1*, and *pDONRP2rP3-mCitrine* were recombined into *pK7m34GW* to generate *pK7m34GW-pBRI1:BRI1^S858A;T872A^-mCit*. The CDS fragments of WT *BRI1* and *BRI1^S858A;T872A^* were amplified using *pDONR221-BRI1* and *pDONR221-BRI1^S858A;T872A^* as templates. The fragments were in-fusion cloned into *pCB302-p35S:GFP* digested with *Bam*HI and *Stu*I to generate *pCB302-p35S:BRI1-GFP* and *pCB302-p35S:BRI1^S858A;T872A^-GFP*. cDNA fragments of *GRF6* and *PUB13* were amplified by using *pET28a-GRF6* (Liu et al., 2017) or Col-0 cDNA as templates. The fragments were in-fusion cloned into *pMDC32-2xp35S:EGFP-HA* digested with *Bam*HI and *Stu*I to generate *pMDC32-2xp35S:GRF6-HA* and *pMDC32-2xp35S:PUB13-HA*, respectively, or were ligated into *pZP211-p35S:3xFlag* digested with *Bam*HI and *Spe*I to generate *pZP211-p35S:GRF6-3xFlag*.

The clones used for protein expression were made as follows: the cDNAs encoding the *BRI1-CD*, *BRI1^S858A;T872A^-CD*, *BRI1-JK*, *BRI1-KC*, and *BRI1-K* were in-fusion cloned into *pGEX4T-1* digested with *Bam*HI and *Sal*I to generate *pGEX4T-GST-BRI1-CD*, *pGEX4T-GST-BRI1^S858A;T872A^-CD*, *pGEX4T-GST-BRI1-JK*, *pGEX4T-GST-BRI1-KC*, and *pGEX4T-GST-BRI1-K*, respectively. *pGEX4T-BRI1^S858A^-CD* and *pGEX4T-BRI1^K911E^-CD* were obtained by site-directed mutagenesis with *pGEX4T-BRI1-CD* as template. The cDNAs encoding the full length of selected *GRFs* and the *BAK1-CD* were in-fusion cloned into *pET28a* digested with *Bam*HI and *Sal*I to generate *pET28a-HIS-GRFs*, and *pET28a-HIS-BAK1-CD*, respectively. *pET28a-HIS-BAK1^K317E^-CD* obtained by site-directed mutagenesis with *pET28a-HIS-BAK1-CD* as template. For the split-ubiquitin based Y2H experiment, full-length *BRI1* and *BRI1^S858A;T872A^* were restriction digested and ligated into *pBT3-STE* and *GRF6* into *pPR3-N* with *Sfi*I.

The *grf1,2,3,4,5,6,7,8* mutant was generated by disrupting *GRF2*, *GRF3*, *GRF5,* and *GRF7* simultaneously in the *grf1,4,6,8* background by means of the CRISPR-Cas9 genome editing technology. For each gene, two gRNAs were designed with the web tool CRISPOR (http://crispor.org) (Concordet and Haeussler, 2018). The eight sgRNAs targeting *GRF2*, *GRF3*, *GRF5*, or *GRF7* were cloned into a single CRISPR/Cas9 vector using the Golden Gate technology as previously described (Decaestecker et al., 2019). Briefly, the gRNA pairs were cloned into the Golden Gate entry modules *pGG-A-AtU6ccdB-B*, *pGG-B-AtU6ccdB-C*, *pGG-C-AtU6ccdB-D,* or *pGG-D-AtU6ccdB-E*. Four-armed entry vectors, each containing two sgRNAs, together with the *pGG-E-linker-G* vector were assembled into a CRISPR destination vector *pFASTRK-pRPS5Ap-AtCas9-NLS-P2A-mCherry-G7T-A-Cm^R^-ccdB-G*.

### Generation of transgenic plants

*Agrobacterium tumefaciens* (strain GV3101)-mediated transformation into the heterozygous *bri1* null mutant, Col-0, *grf9,10,11^RNAi^*, *grf1,4,6,8*, *grf1,2,3,4,5,6,7,8*, *pBRI1:BRI1-mCit/bri1*, or *pBRI1:BRI1^S858A;T872A^-mCit/bri1* was used to create stable transgenic lines. The transformants were screened by germination on 0.6% (w/v) agar plates (without sucrose and with 100 mg/L timentin) containing 50 mg/L kanamycin, 10 mg/L BASTA, or 25 mg/L hygromycin according to the plant resistance marker in the binary vectors.

### Chemical treatments

MG-132 (Cell Signaling Technology; 10 mM stock in DMSO), Brassinolide (Cayman; 10 mM stock in DMSO for co-IP assays, or TCI Europe N.V.; 20 mM stock in DMSO for hypocotyl growth and BES1 dephosphorylation assays), Brassinazole (Macklin; 10 mM stock in DMSO), Brefeldin A (Psaitong; 50 mM stock in DMSO), CHX (Merck; 50 mM stock in DMSO), and Concanamycin A (Cayman; 115 µM stock in acetonitrile) were used at the concentrations indicated and plants were treated for the indicated periods.

### Recombinant protein expression, purification, and GST pull-down

Fusion proteins were generated from bacterial protein expression vectors as described above in *Escherichia coli* BL21 or Rosetta strain grown in Luria-Bertani medium supplemented with 0.5 mM isopropyl β-D-1-thiogalactopyranoside and induced for 16 h at 16°C. The GST- or 6xHIS-fused proteins were purified with glutathione sepharose 4B GST-tagged protein purification resin (Solarbio) or His-tag protein purification kit (CWBIO), respectively, according to the manufacturers’ standard procedures.

Approximately 10 µg of GST or GST-fused proteins in the pull-down buffer (20 mM Tris-HCl, pH 7.5, 100 mM NaCl, 0.1 mM EDTA, 0.2% [v/v] Triton X-100) were incubated with prewashed 20 µL glutathione agarose beads (Solarbio) at 4°C for 2 h with gentle shaking. Beads were washed with 400 µL pull-down buffer once, followed by incubation with 10 µg of 6xHIS-fused proteins by shaking at 4°C for another 1 h. Afterward, the beads were washed four times with 400 µL washing buffer (20 mM Tris-HCl, pH 7.5, 300 mM NaCl, 0.1 mM EDTA, 0.5% [v/v] Triton X-100). Beads were boiled in 50 µL of 2×SDS protein loading buffer for 5 min and detected by immunoblotting (IB) with a-GST (TRANS, X015, 1:3,000) and a-HIS (TRANS, E065, 1:3,000) antibodies. For blocking and antibody dilutions, 5% (w/v) nonfat milk powder in 0.1% (v/v) Tween-20 containing Tris-buffered saline (TBST) was used.

### Immunoblot analysis and co-immunoprecipitation

For autophosphorylation analysis, equal amounts of recombinant GST-tagged wild-type, S858A;T872A-mutated, S858A-mutated, and kinase-dead (BRI1^K911E^) BRI1-CD proteins were loaded and detected by IB with α-pS/T (ECM Biosciences, PP2551, 1:1,000), α-pY (Invitrogen, 10-5001-82, 1:3,000), α-pS_858_ (Oh et al., 2012) and a-GST antibodies. For *in vitro* kinase assay, 1 µg of GST-BRI1-CD was incubated with 10 µg HIS-tagged kinase-dead BAK1-CD (HIS-BAK1^K317E^-CD) with or without the presence of 2 µg HIS-GRF6 in kinase reaction buffer (10 mM Tris-HCl, pH 7.5, 5 mM MgCl_2_, 2.5 mM EDTA, 50 mM NaCl, 0.5 mM dithiothreitol [DTT] and 50 µM ATP) at the final volume of 20 µL. After gentle shaking for 2 h at room temperature, samples were denatured by adding 5 µL 5×SDS loading buffer and boiled for 5 min. The samples were detected by IB with a-pS/T, a-GST, and a-HIS antibodies.

For the BRI1 and BAK1 expression and BES1 dephosphorylation assays, 5-day-old seedlings untreated or treated with 50 µM MG-132 or BL (0, 1, and 10 nM) were pulverized and extracted with 2xSDS loading buffer and directly boiled for 10 min, followed by centrifugation at 13,000*g* for 10 min. The supernatants were detected by IB with α-GFP (TRANS, D044, 1:3,000), α-BRI1 (Agrisera, AS12 1859, 1:5,000), α-BAK1 (Agrisera, AS12 1858, 1:5,000), α-BES1 (Yin et al., 2002), β-ACTIN (CWBIO, CW0264, 1:10,000), and α-tubulin (Sigma-Aldrich, T5168, 1:10,000) antibodies. For BES1 dephosphorylation assays, the ratio of the dephosphorylated BES1 to the total BES1 was quantified based on the band intensity using ImageJ.

For co-IP in *N. benthamiana*, *A. tumefaciens* (strain LBA4404) expressing *p35S:BRI1-HA*, *p35S:GFP* or *p35S:GRF-GFP* for each of the 7 *GRFs* selected were resuspended in infiltration buffer (10 mM MgCl_2_, 10 mM MES pH 5.6, 0.1 mM acetosyringone) for 2 h at room temperature. *p35S:BRI1-HA* culture was mixed with *p35S:GFP* or each of the *p35S:GRF-GFP* to a final OD_600_=0.5. Two days after infiltration into *N. benthamiana*, leaves were harvested, ground to powder with liquid nitrogen, and solubilized in extraction buffer [50 mM Tris-HCl, pH 7.5, 150 mM NaCl, 2.5 mM EDTA, 10% (v/v) glycerol, 1% (v/v) NP-40, 5 mM DTT, complete EDTA-free protease inhibitor cocktail (SIGMA)] in a 1:2 (w:v) for 30 min at 4°C with gentle shaking. The extracts were centrifuged twice at 16,000*g* at 4°C for 20 min and the supernatants were incubated with α-GFP-coated magnetic beads (AlpaLifeBio) or α-HA-coated magnetic beads (AlpaLifeBio) for 2 h at 4°C, followed by washing three times with 1 mL of washing buffer (same as extraction buffer but without protease inhibitor). Beads were eluted with 40 µL 2xSDS loading buffer and boiled for 10 min. Samples were analyzed by IB with α-GFP and α-HA (Abmart, M20003, 1:3,000) antibodies.

Arabidopsis microsomal fractions were isolated according to the published protocol (Liu et al., 2020) with the minor modification that seedlings were treated with only 50 µM MG-132 for 5 h. For co-IP with Arabidopsis rosette leaves as material, the same protocol was used as described above for co-IP in *N. benthamiana*. For co-IP assays testing the dependency of BRs level, 5-day-old Arabidopsis seedlings were first treated with 2 μM BRZ for 3 days, followed by 1 μM BL treatment or continuously with 2 μM BRZ for 1.5 h. The seedlings were harvested for co-IP with α-GFP-coated magnetic beads as above and blotted by the corresponding antibodies. For solubilized microsomal fractions, the same co-IP protocol was used as above, starting from incubation with the prewashed α-GFP-coated magnetic beads. Samples were analyzed by IB with α-GFP, α-HA, α-ubiquitin (Cell Signaling Technology, 3933S, 1:2,000), and α-Flag (TRANS, HT201, 1:3,000) antibodies.

For blocking and antibody dilutions, 5% (w/v) BSA powder in 0.1% (v/v) TBST was used for α-pS/T, α-pY, and α-pS_858_ IBs, and TBST-containing 5% (w/v) nonfat milk powder for the remaining IBs.

### qRT-PCR

Total RNA was extracted from 5-day-old seedlings with Ultrapure RNA kit (CWBIO). cDNA was synthesized with the EasyScript One-Step gDNA Removal and cDNA Synthesis SuperMix (TRANS). qRT-PCRs were run with TransStart Tip Green qPCR SuperMix Tip Green (TRANS) on a QuantStudio 6 Flex (Applied Biosystems, Thermo Fisher Scientific). The *BRI1* and *GRF* expressions were normalized to that of *ACTIN2*. The cycling conditions were as follows: 50°C, 2 min, and 95°C, 1 min (pre-incubation); 95°C, 15 s, and 60°C, 1 min (40 cycles of amplification); 95°C, 15 s, 60°C, 1 min, and 95°C, 15 s (melting curve).

### Split-ubiquitin based Y2H

Constructs used for the Y2H assay were generated as described above. Plasmids with different combinations were co-transformed into the yeast strain NMY51 by the LiAc-mediated yeast transformation (Gietz and Schiestl, 2007). Positive yeast transformants were screened on SD/-Trp-Leu medium, and protein interactions were selected on SD/-Trp-Leu-His-Ade medium.

### Confocal imaging and data analysis

For analysis of BRI1-GRF6 rBiFC, BRI1 internalization and vacuole targeting, *N. benthamiana* leaves or the meristem zone of Arabidopsis roots were imaged with a 20x objective lens (numerical aperture, 0.8) or a 60x oil corrected immersion objective lens (numerical aperture, 1.4) on a confocal laser scanning microscope (LSM880; Zeiss). The excitation/emission wavelengths used were 488 nm/505-552 nm for GFP (or YFP) and 594 nm/605-639 nm for RFP. Images were converted to 8-bit in ImageJ for YFP and RFP intensity quantification in rBiFC assays, and for BRI1 (or BRI1^S858A;T872A^)-GFP fluorescence signal intensity measurements. Regions of interest were selected based on the PM or cytosol localization. The averages of the top bright 10% pixels from the whole PM or intracellular part were used to calculate the YFP/RFP ratios, or the relative PM/intracellular fluorescent intensity as previously described (Liu et al., 2020). BFA body size was measured as previously described (Luo et al., 2015).

### AlphaFold structure prediction and structure illustration

The structural configurations of BRI1-CD/GRF6, BRI1^S858/T872-P^-CD/GRF6, BRI1-CD/BAK1-CD and BRI1-CD/BAK1-CD/GRF6 were predict using AlphaFold ([AlphaFold Server](https://alphafoldserver.com/)), a dedicated online server for protein three-dimentional structure prediction (Jumper et al., 2021). The structures were visualized by PyMOL (http://www.pymol.org/pymol).

### Quantification and Statistical Analysis

Statistical analyses were carried out in Excel with build-in formulas or XL Toolbox NG. The *P* values were calculated with two-tailed Student’s unpaired *t*-test analysis for binary comparison, or with one-way ANOVA and Tukey’s post hoc honestly significance test for comparisons of more than two genotypes or combinations. The measurements shown in box plots display the first and third quartiles and are split by medians (center lines), with whiskers extending to 1.5-fold the interquartile range from the 25th and 75th percentiles. Outliers are represented by dots. The box plots were generated at BoxPlotR (shiny.chemgrid.org/boxplotr/). The measurements shown in graphs display the averages and standard errors.

### Accession Numbers

Sequence data from this article can be found in the GenBank/EMBL data libraries under accession numbers: BRI1 (At4G39400); GRF1 (AT4G09000); GRF2 (AT1G78300); GRF3 (AT5G38480); GRF4 (AT1G35160); GRF5 (AT5G16050); GRF6 (AT5G10450); GRF7 (AT3G02520); GRF8 (AT5G65430); GRF9 (AT2G42590); GRF10 (AT1G22300); GRF11 (AT1G34760); PUB13 (AT3G46510); BAK1 (AT4G33430); BKI1 (AT5G42750); and BES1 (AT1G19350).

## Author contributions

D.Liu, E.R., and D.Luo initiated the project and designed experiments. L.L. generated most of the plant materials, most of the clones, the CRISPR constructs, the *grf1,2,3,4,5,6,7,8* mutant, and performed BRI1 ubiquitination assay, part of the kinase assays and BRI1 stability assays. C.Yin performed part of the co-IP assays, most of the Y2H assays, and part of GST pull-down assays and most of the *in vitro* kinase assays. C.Yan performed part of the Y2H experiments, all the qRT-PCR analysis, most of the genotyping and protein expression assays, and performed part of the co-IP assays and GST pull-down assays. Z.H performed BL treated growth and BES1 dephosphorylation assays. I.V. generated part of the plant materials. Y.L. and G.X. participated in the genotyping, co-IP assays, and part of the cloning work. B.T. performed the BAK1 stability assay, and part of protein purification. X.F. and W.C. participated in protein expression, purification, and GST pull-down assays. D.Liu performed the confocal imaging and data analysis. Y.W. gave guidance on the writing and revision of the manuscript. D.Luo, E.R. and D.Liu wrote the manuscript. All authors revised the manuscript.

## Supplementary Data

**Supplementary Figure S1.**
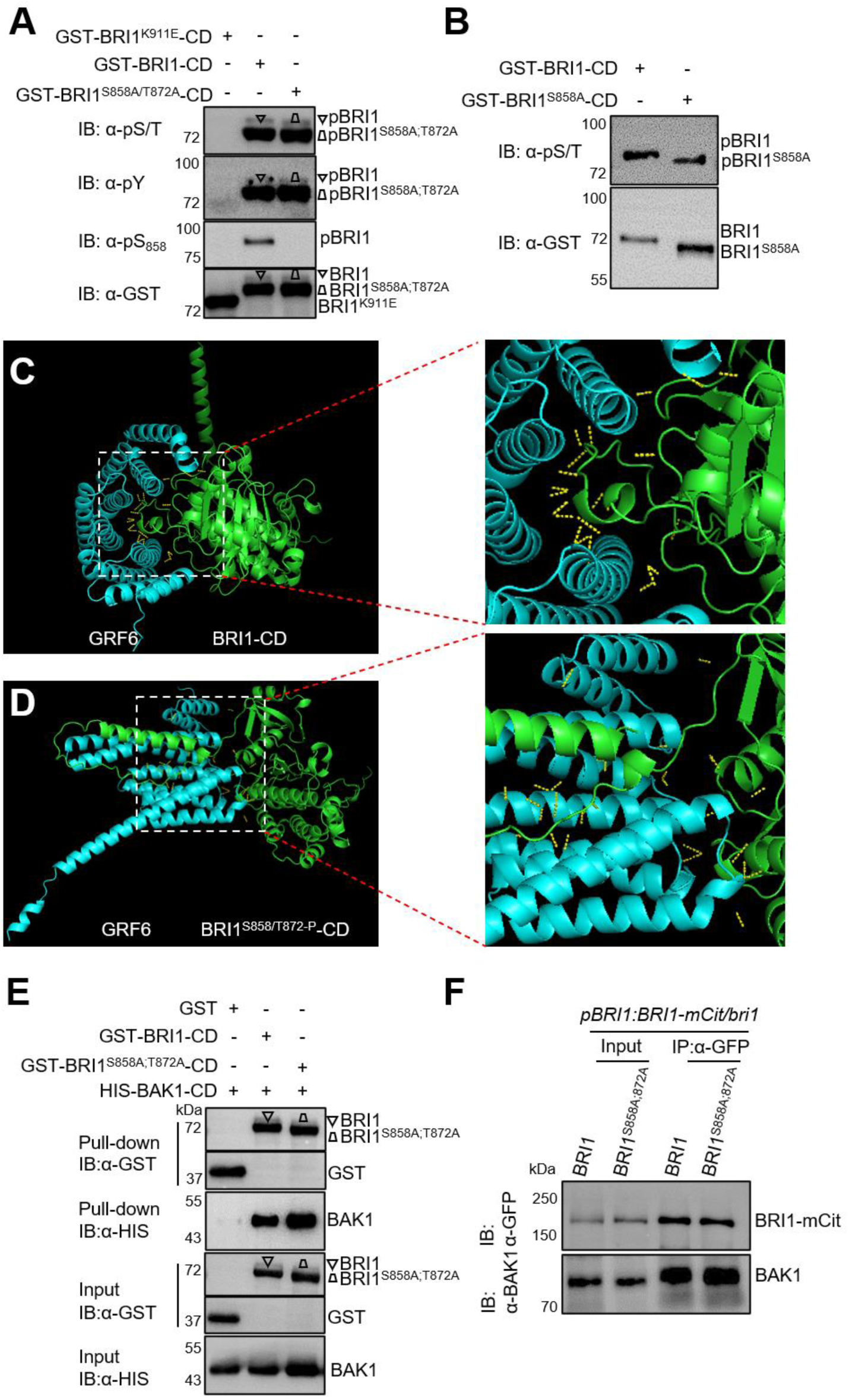
Effect of S858A and T872A mutations on BRI1 kinase activity and BRI1-BAK1 association. **A** and **B**) Effects of S858A;T872A double or S858A single residue substitutions in BRI1-CD on their autophosphorylation status. Equal amounts of recombinant N-terminally GST-tagged cytoplasmic domain (CD) of wild-type BRI1, kinase-dead BRI1^K911E^, BRI1^S858A^ and BRI1^S858A;T872A^ BRI1 mutants were used *in vitro* kinase reaction, followed by immunoblotting (IB) with modification-specific antibodies (α-pS/T, α-pY, and α-pS_858_). pBRI1-CD and pBRI1^S858A;T872A^-CD indicate the specifically phosphorylated proteins. Different shapes indicate GST-BRI1-CD or GST-BRI1^S858A;T872A^-CD, or phosphorylated ones, respectively. Protein loading is shown by IB assay with an α-GST antibody. **C** and **D**) Three-dimensional structure of BRI1-CD/GRF6 dimmer and BRI1^S858/T872-P^-CD/GRF6 dimmer predited by AlphaFold. Wild-type BRI1-CD or BRI1-CD with S858 and T872 phosphorylated (BRI1^S858/T872-P^-CD), together with GRF6 protein sequences were entried at AlphaFold Server ([AlphaFold Server](https://alphafoldserver.com/)) to generate the BRI1-CD/GRF6 dimmer (**C**) or BRI1-CD/BRI1^S858/T872-P^-CD dimmer (**D**) structures, repectively. BRI1-CD or BRI1^S858/T872-P^-CD, and GRF6 are colored in green and cyan, respectlvely. The polar contacts between chains predicted by PyMOL are indicated with dashed yellow lines. The regions forming polar contacts were enlarged and shown on the right. **E** and **F**) S858A and T872A mutations in BRI1 on BRI1-BAK1 association. **E**) BAK1 associations with wild-type BRI1 and BRI1^S858A;T872A^ mutant determined by *in vitro* GST pull-down. HIS-fused BAK1 cytoplasmic domain (BAK1-CD) was incubated with free GST, GST-fused BRI1-CD, or BRI1^S858A;T872A^-CD. The beads were collected and washed, and the pull-down fractions were followed by IBs with α-GST and α-HIS antibodies. Protein inputs were also determined by α-GST and α-HIS IBs. **F**) BAK1 associations with wild-type BRI1 and BRI1^S858A;T872A^ mutant in Arabidopsis. Eight-day-old Arabidopsis transgenic plants expressing mCit-tagged BRI1 or BRI1BRI1^S858A;T872A^ were first treated with BRZ at 2 µM for 3 days, followed by a treatment with 1 µM BL for 90 min, and co-IP assays using α-GFP-coated beads were performed with proteins isolated from these samples. The proteins were subjected to IB using α-BAK1 and α-GFP antibodies.

**Supplementary Figure S2.**
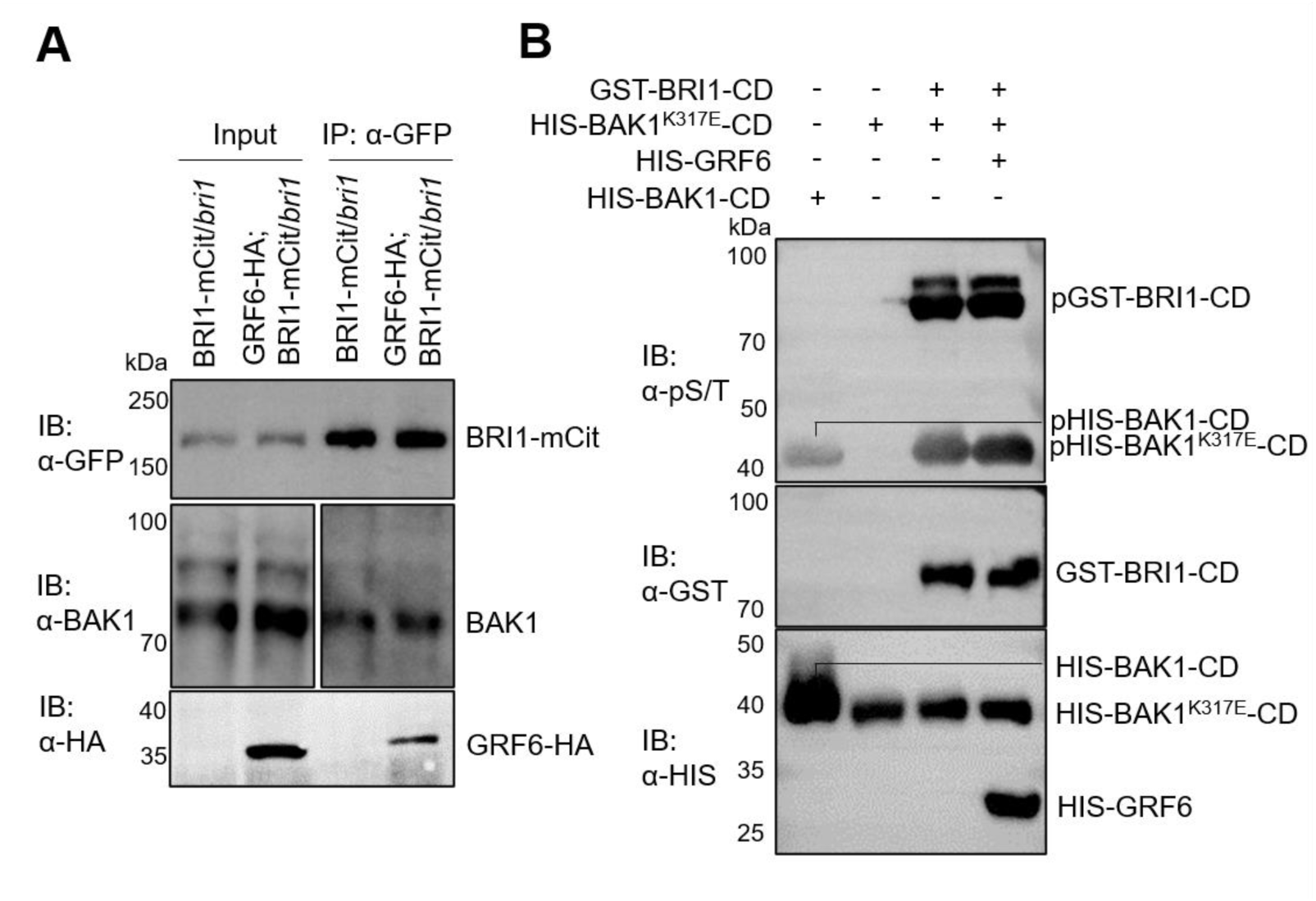
BRI1 association with BAK1 and BRI1 kinase activity were unaffected by the presence of GRF6. **A**) The association between BRI1 and BAK1 in Arabidopsis was unaffected by GRF6. Eight-day-old Arabidopsis transgenic plants expressing BRI1-mCitrine (mCit) alone or together with GRF6-HA were first treated with 2 µM BRZ for 3 days followed by 1 µM BL for 90 min, and subjected to co-IP using α-GFP-coated beads. The proteins were detected by IB with α-GFP, α-BAK1 and α-HA antibodies. **B**) Effect of GRF6 binding on BRI1 kinase activity. Recombinant kinase-dead BAK1 (HIS-BAK1^K317E^-CD), HIS-GRF6, and GST-BRI1-CD proteins were incubated alone or in combination, as indicated. BRI1 or BAK1 phosphorylation was detected by IB with α-pS/T, and protein loading was determined by IB using α-GST and α-HIS antibodies.

**Supplementary Figure S3.**
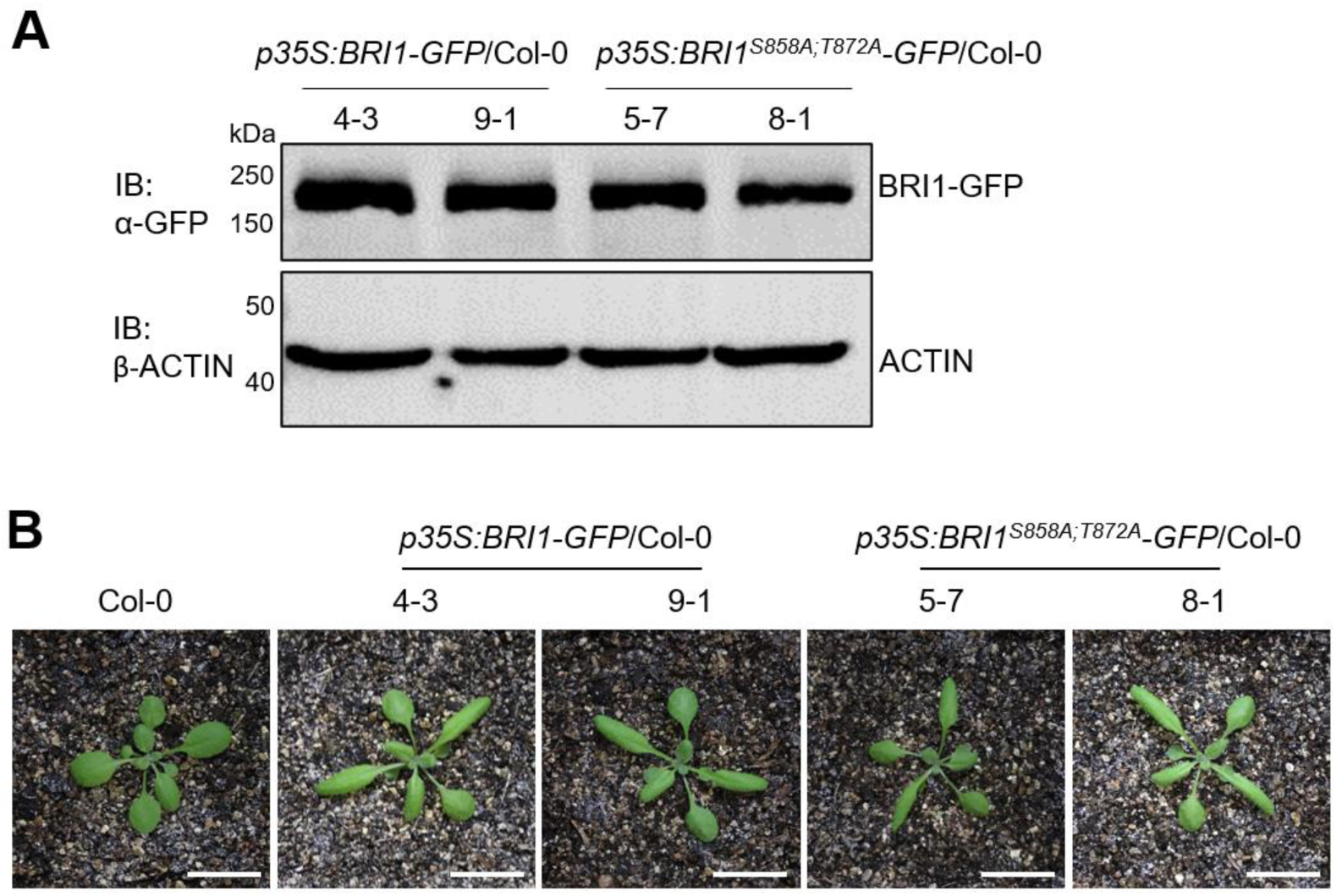
Characterization of *p35S:BRI1-GFP*/Col-0 and *p35S:BRI1^S858A;T872A^-GFP*/Col-0 transgenic plants. **A**) IB analysis of BRI1 expression. Total proteins were isolated from 5-day-old seedlings of the transgenic lines and detected using an α-GFP antibody (top). ACTIN was used as loading control by IB assay with the β-ACTIN antibody (bottom). **B**) Growth phenotypes of soil-grown 26-day-old transgenic lines. Scale bar, 1 cm.

**Supplementary Figure S4.**
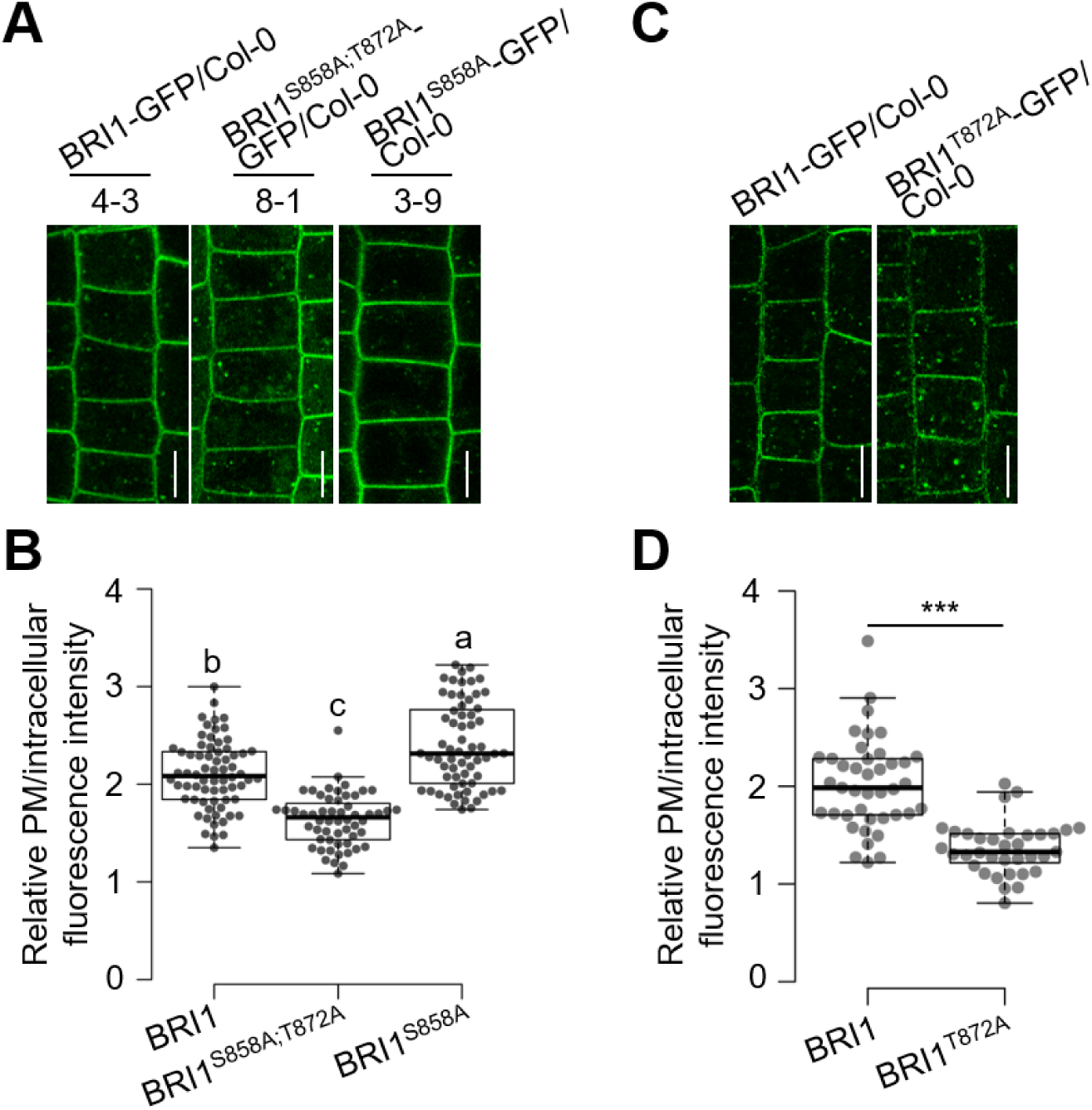
Internalization behaviors of BRI1^S858A^ and BRI1^T872A^. **A** and **C**) Images of epidermal cells from root meristems of 5-day-old homozygous (**A**) and 7-day-old screened T1 (**C**) Arabidopsis seedlings expressing different versions of BRI1 tagged with GFP pretreated with CHX at 50 µM for 1.5 h. **B and D**) Measurements of the relative PM versus intracellular BRI1-GFP fluorescence intensity derived from the images displayed in the upper panel. At least 30 cells from four seedlings were measured for each background. Different letters indicate a significant difference with others (*P* < 0.05, one-way ANOVA followed by Tukey’s test). \*\*\**P* < 0.001, Student *t*-test. Scale bars, 10 µm.

**Supplementary Figure S5.**
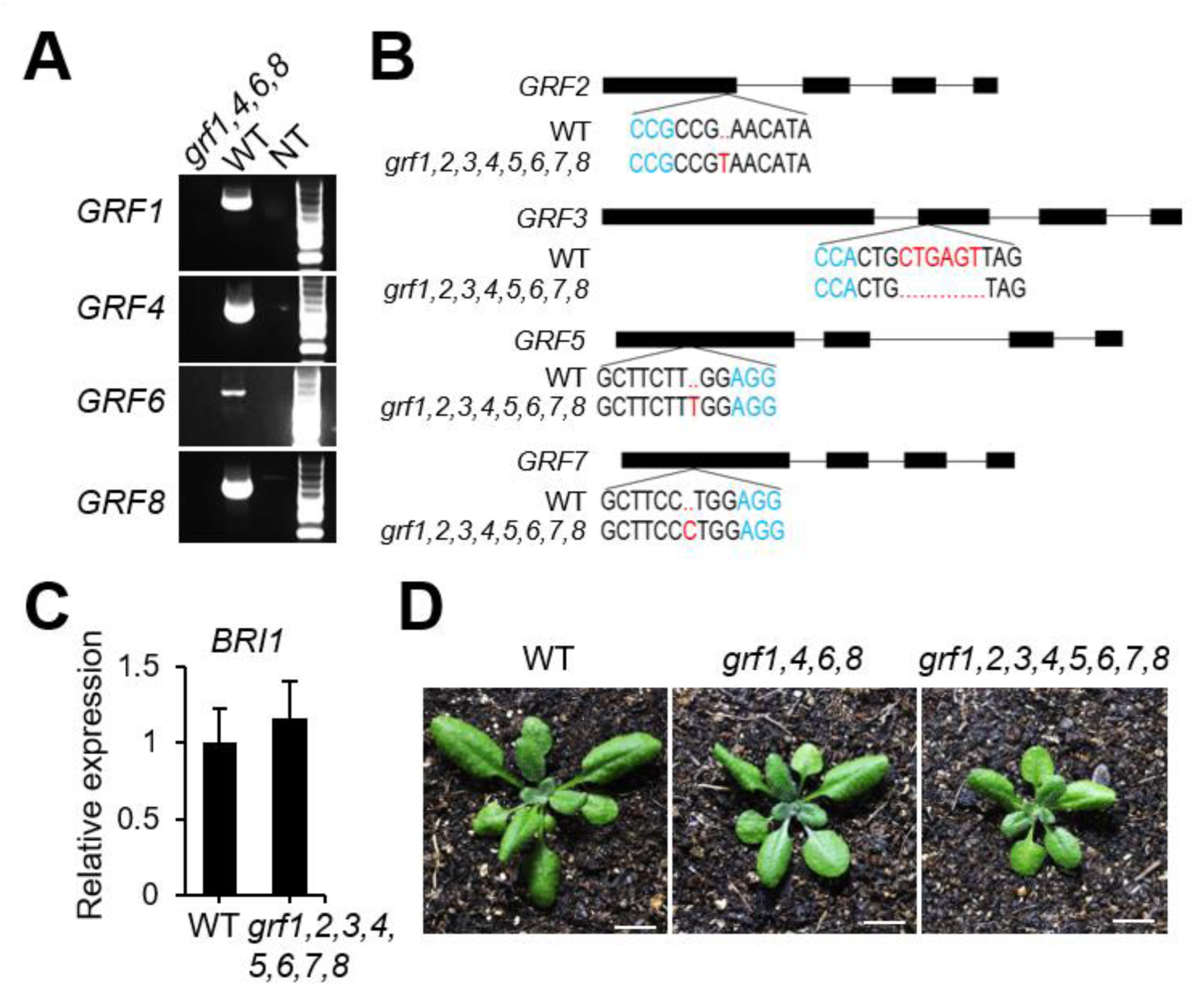
Molecular characterization of the quadruple *grf1,4,6,8* mutant and octuple *grf1,2,3,4,5,6,7,8* mutant. **A**) Genotyping of quadruple *grf1,4,6,8* mutant. Genomic DNA isolated from *grf1,4,6,8* mutant and wild-type plants and the negative control (NT) were used for PCR analysis with gene-specific primers. **B**) Generation of *CR-2,3,5,7* quadruple mutants in *klpc* background using CRISPR-Cas9. The gene structures and resulting mutations in *GRF2*, *GRF3*, *GRF5*, and *GRF7* are illustrated. **C**) qRT-PCR analyses of *BRI1* (*n* =3, *n*, biological replicates). Total RNA was isolated from 7-day-old seedlings. *ACTIN2* expression was used as an internal control. Error bars indicate standard deviation (SD). **D**) Growth phenotypes of soil-grown indicated lines at 22°C, 56% relative humidity, and 16-h light/8-h dark for 26 days. Scale bar = 1 cm.

**Supplementary Figure S6.**
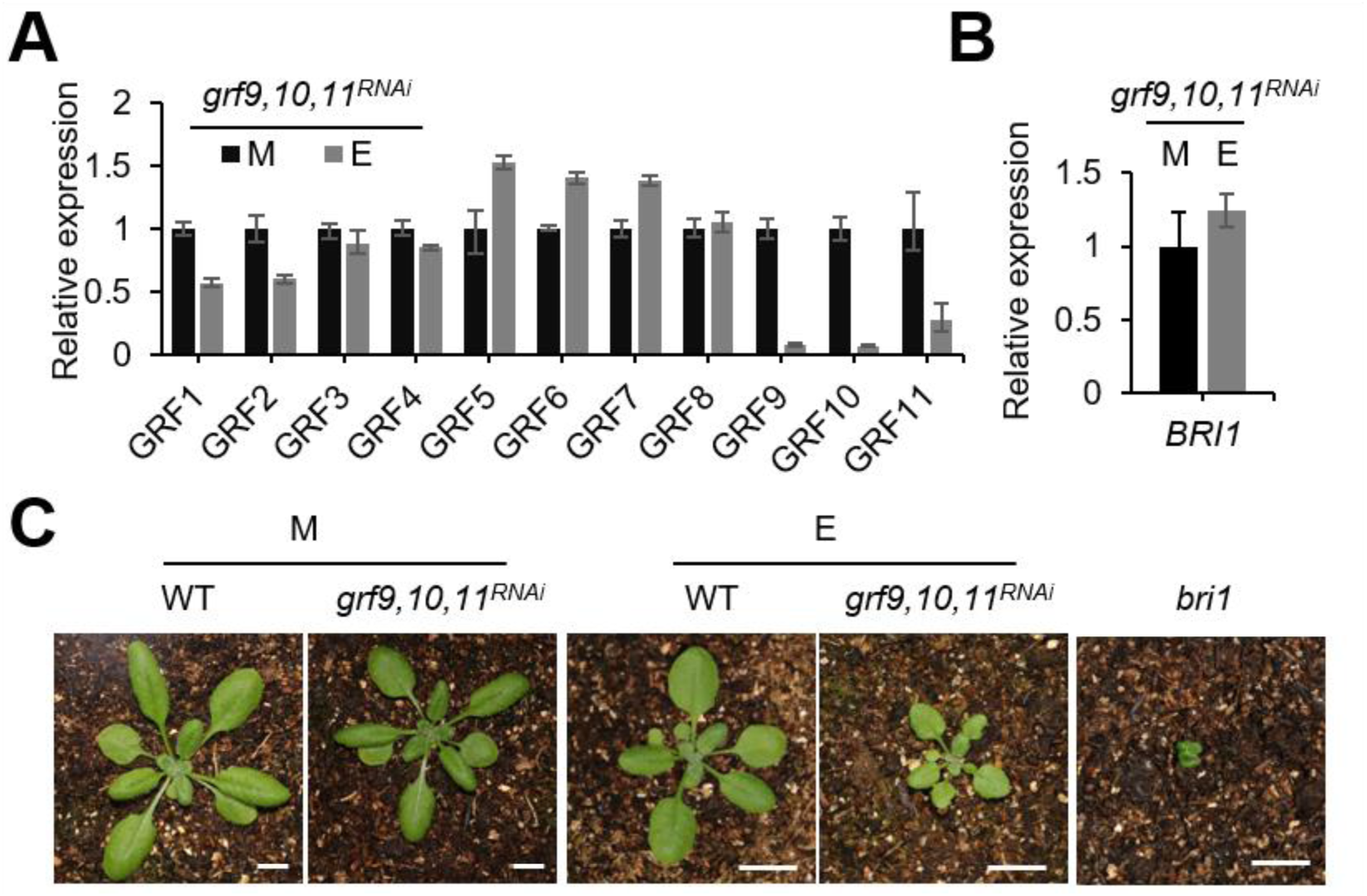
Molecular characterization of the ethanol-inducible amiRNA line, *grf9,10,11^RNAi^*. **A** and **B**) qRT-PCR analyses of *GRF1-11* and *BRI1* (*n* =3, *n*, biological replicates). Total RNA was isolated from 5-day-old seedlings grown in a medium with (E) or without (M) 0.1% (v/v) ethanol. *ACTIN2* expression was used as an internal control. Error bars indicate standard deviation (SD). **C**) Growth phenotypes of soil-grown 22-day-old plants. The seeds were sown directly into the soil. Plants were sprayed with H_2_O or 1% (v/v) ethanol-containing water from day 7 every other day (two spays per plant) until day 21. Plants were photographed on day 22. Scale bar in (**C**), 1 cm.

**Supplementary Figure S7.**
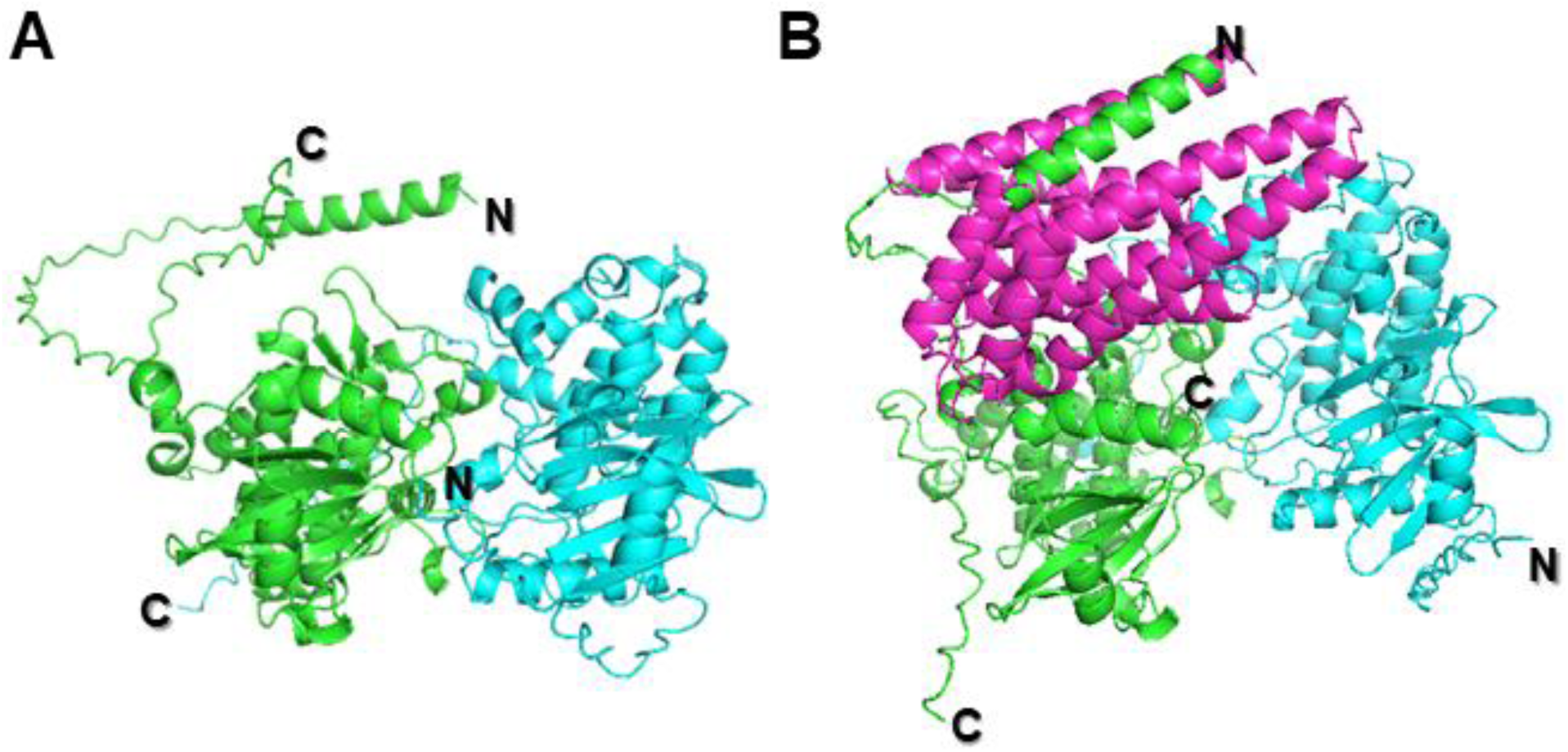
Three-dimensional structure of BRI1-CD/BAK1-CD dimmer and BRI1-CD/BAK1-CD/GRF6 trimer predited by AlphaFold. BRI1-CD, BAK1-CD, together without or with GRF6 protein sequences were entried at AlphaFold Server ([AlphaFold Server](https://alphafoldserver.com/)) to generate the BRI1-CD/BAK1-CD dimmer (**A**) or BRI1-CD/BAK1-CD/GRF6 trimer (**B**) structures, repectively. BRI1-CD, BAK1-CD, and GRF6 are colored in green, cyan, and magenta, respectlvely. The N’ and C’-terminals of BRI1-CD and BAK1-CD were labeled with N and C. The figures were generated with PyMOL.

**Supplementary Table S1**. Primers used in this study.

## Acknowledgments

We thank Michael Hothorn (University of Geneva, Geneva, Switzerland) for discussions of the project, Yanhai Yin (Iowa State University, Ames, USA), Shuhua Yang (China Agricultural University, Beijing, China), Claudia Oecking (University of Tübingen, Tübingen, Germany), Han Jiang (Shandong Agricultural University, Tai’an, China), Xuelu Wang (Henan University, Kaifeng, China), Shujing Wu (Shandong Agricultural University, Tai’an, China), and Steve Huber (University of Illinois, Urbana, USA) for providing the α-BES1 antibody, the *pET28a-GRF6* construct, the amiRNA line *grf9,10,11^RNAi^*, the split-ubiquitin system, the *35S::BKI1-GFP*/Col-0 and *35S::BES1-S-GFP*/Col-0 lines, the *pCB302-35S:GFP* constructs, and α-pS_858_ antibody, respectively, and Martine De Cock (VIB-Ghent University, Ghent, Belgium) for help in preparing the article. HOME for Researchers is acknowledged for linguistic assistance during the preparation of this manuscript.

## Data availability

All the materials used in this study is available upon request to the corresponding authors.

